# Genomic loci susceptible to systematic sequencing bias in clinical whole genomes

**DOI:** 10.1101/679423

**Authors:** Timothy M. Freeman, Genomics England Research Consortium, Dennis Wang, Jason Harris

## Abstract

Accurate massively parallel sequencing (MPS) of genetic variants is key to many areas of science and medicine, such as cataloguing population genetic variation and diagnosing genetic diseases. Certain genomic positions can be prone to higher rates of systematic sequencing and alignment bias that limit accuracy, resulting in false positive variant calls. Current standard practices to differentiate between loci that can and cannot be sequenced with high confidence utilise consensus between different sequencing methods as a proxy for sequencing confidence. These practices have significant limitations and alternative methods are required to overcome these.

We have developed a novel statistical method based on summarising sequenced reads from whole genome clinical samples and cataloguing them in “Incremental Databases” that maintain individual confidentiality. Allele statistics were catalogued for each genomic position that consistently showed systematic biases with the corresponding MPS sequencing pipeline. We found systematic biases present at ∼1-3% of the human autosomal genome across five patient cohorts. We identified which genomic regions were more or less prone to systematic biases, including large homopolymer flanks (odds ratio=23.29-33.69) and the NIST high confidence genomic regions (odds ratio=0.154-0.191). We confirmed our predictions on a gold-standard reference genome and showed that these systematic biases can lead to suspect variant calls within clinical panels.

Our results recommend increased caution to address systematic biases in whole genome sequencing and alignment. This study provides the implementation of a simple statistical approach to enhance quality control of clinically sequenced samples by flagging variants at suspect loci for further analysis or exclusion.

## Introduction

DNA sequencing is an imperfect process, and although error rates are low, mistakes in identifying genomic variants can still occur. While the sources of random sequencing errors are relatively well understood (Ma et al. 2019; Benjamini and Speed 2012), identifying systematic errors in whole genomes sequenced in a clinical or commercial setting is not always possible due to restrictions in gathering information about the samples and sequencing processes. These errors could cause incorrect decisions on the presence or absence of disease relevant variants in the genome and influence clinical and research decisions (Goldfeder et al. 2016).

One of the major challenges to improving variant detection is that certain regions of the genome are prone to higher rates of systematic sequencing or alignment errors, which can result in the false identification of variants at low allelic fraction. In the case of diploid genotype calls, variants are expected to be around 50% or 100% allelic fraction, corresponding to heterozygous and homozygous loci. However, real variants sometimes occur at low allelic fractions, such as somatic variants in tumours and in cases of mosaicism, where nearby cells sampled together can show genetic heterogeneity within the sample (King et al. 2017; Vattathil and Scheet 2016). In these cases the ability to identify loci that systematically exhibit a low allelic fraction across individuals becomes critical, since these artifacts may be mis-identified as variant alleles.

Lists of ‘high confidence’ loci from gold-standard reference genomes are sometimes used for quality control purposes in clinical and commercial sequencing laboratories, since they leave out regions which cannot be sequenced reliably by any technology, although they are not designed to reflect high sequencing accuracy. For example, The National Institute of Standards and Technology Genome in a Bottle consortium (NIST GIAB) has proposed a list of ‘high confidence’ genomic regions to be used for benchmarking different sequencing methods, developed using a top-down approach, by analysing the consensus between different sequencing technologies and variant callers for the same genomic samples to develop a ‘truth set’ of variant calls (Zook et al. 2014, 2019). However, because genomic regions only require at least one sequencing method showing no evidence of systematic error to be included in the ‘high confidence’ list, the sequencing pipeline being used by any one scientist could be a different method that is affected by systematic error, so filtering out genomic regions not in the ‘high confidence’ list does not guarantee high sequencing accuracy in all remaining regions. This disparity between the ‘high confidence’ set and regions with high sequencing accuracy for any one sequencing pipeline is likely to increase over time as more genomic regions are included in the ‘high confidence’ set which can be sequenced by long and linked-read based methods, but which show systematic biases with short-read based sequencing. Another drawback to this is that clinically collected samples can vary in quality and contamination may introduce variants with low allelic fractions not seen in reference genomes. Furthermore, the number of reference genomes used may be quite small, so sample-specific structural variants, which are not representative of the diversity of clinically-sequenced genomes, can cause genomic regions to be missing from the ‘high confidence’ list despite having accurate sequencing for most samples.

Other top-down approaches of evaluating thresholds for allelic fraction or read quality may differ depending on the variant calling pipelines used (Sandmann et al. 2017). Benchmarking these different approaches on cohorts of genomes may be insightful for research, but impractical for clinical applications and it risks leaking sensitive genetic information. Furthermore, standard quality control measures for variant calling can often be overly simplistic, such as fixed read depth thresholds for calling variants across the genome, which are not tailored to wide regional differences in systematic biases. A reliance on high read depth for accurate variant calling increases the costs of sequencing studies, which are forced to compromise between the number of genomes sequenced and the depth of coverage achieved (The 1000 Genomes Project Consortium 2010).

A ‘bottom-up’ approach to learning about sequencing error looks at different cohorts of clinically sequenced genomes independently and does not rely on consensus between multiple sequencing technologies. This study aims to address the limitations affecting other quality control methods described above, by developing a ‘bottom-up’ method to evaluate position-dependent systematic bias in detected allele fractions across whole genomes. We also seek to quantify its utility using results from its application to five small WGS non-cancer cohorts, to detect how common these systematic biases are, the extent to which they affect different genomic regions and to gauge whether the systematic bias predictions from our method are supported by gold-standard reference data.

## Results

### Biased variant allelic fractions occur across the genome and are persistent across individuals

Five separate ethnically-mixed patient cohorts in the USA and UK (**Table 1**) were previously sequenced under confidential conditions using distinct WGS pipelines prior to this study, by Personalis Inc. and the 100,000 Genomes Project. Incremental Databases (IncDBs) were generated using this data, containing between-patient standard deviation of allelic fractions for each autosomal genomic locus, and also the corresponding aggregate allelic fraction values (**Fig. 1**), without maintaining any sample-specific information. This ensured that individuals remained anonymous during the quality control process of examining reads using an IncDB. Aggregate allelic fraction was defined as the mean of all of the allelic fractions across the patient cohort, for that specific allele at a specific single-nucleotide genomic position. All patients were non-cancer patients, and in addition patients in data sets 2-5 all had neurological disorders. The main differences between data sets were the use of the reference genome GRCh37 being used for alignment in data sets 1 and 2, while GRCh38 was used for data sets 3-5, and the use of BWA-MEM (Li 2013) for alignment in data set 1, while the Isaac-aligner (Raczy et al. 2013) was used in data sets 2-5. Since clinical sequencing facilities will continually upgrade their sequencing pipelines with new versions of algorithms and genome builds as time goes on, we wanted to make comparisons between different sequencing protocols in different cohorts of patients. In order to verify our findings, we also sequenced two additional cohorts (data sets 4 and 5) using exactly the same protocol as data set 3.

**Table 1:** Overview of cohorts whole genome sequenced and analysed.

| Data set | Source | Individual genomes | Genome build | Sequencing | Alignment |
| --- | --- | --- | --- | --- | --- |
| **1** | Personalis Inc. (USA) | 150 | GRCh37 | Illumina HiSeqX | BWA-MEM 0.7.12 |
| **2** | 100,000 Genomes Project (UK) | 215 | GRCh37 | Illumina HiSeqX | Isaac-aligner (SAAC00776.1 5.01.27) |
| 3-5 | 100,000 Genomes Project (UK) | 215 | GRCh38 | Illumina HiSeqX | Isaac-aligner (iSAAC-03.16.02.19) |

**Figure 1:**
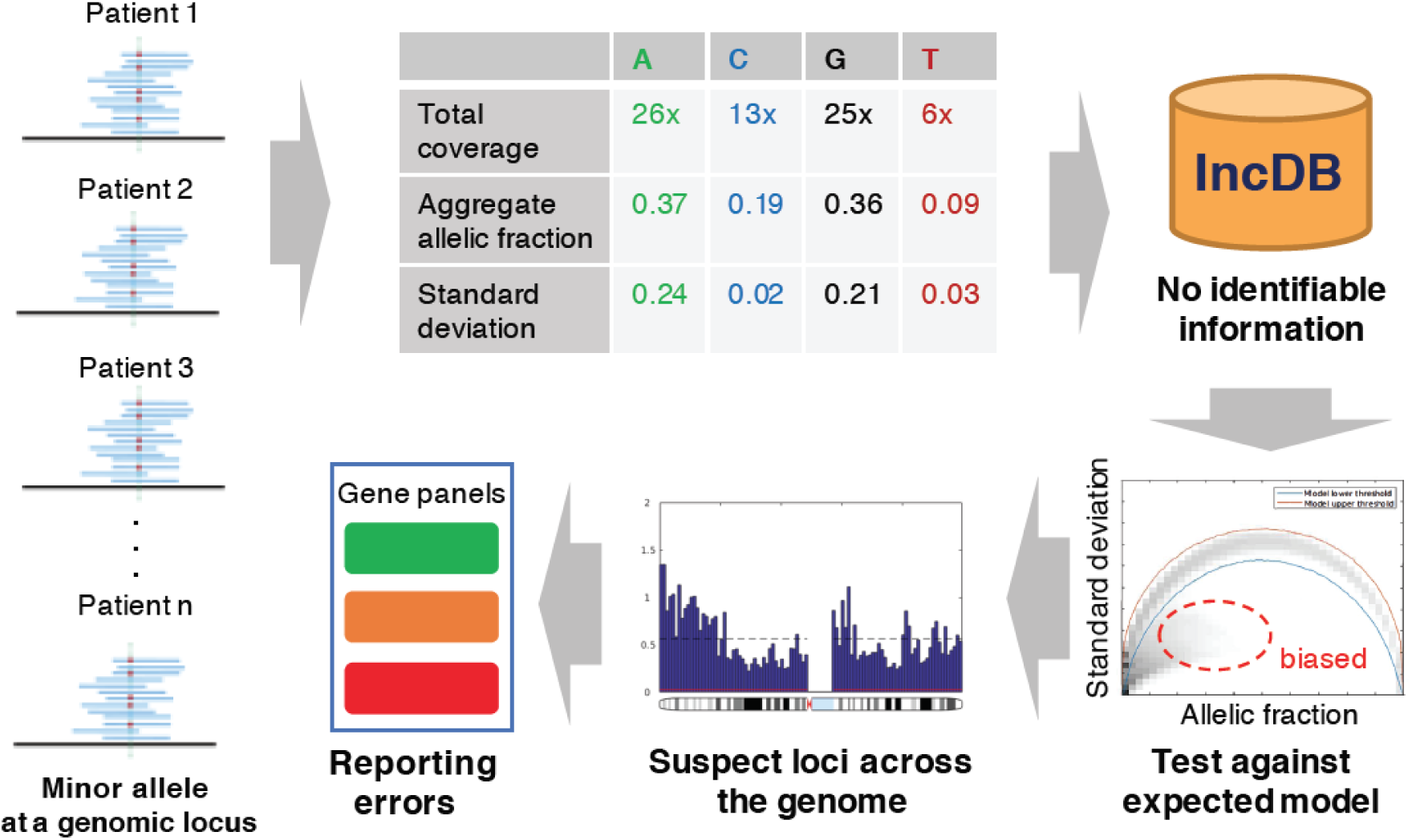
Approach for detecting loci with systematic sequence bias: Alternative allele fractions are collected from a cohort of individuals at every locus and aggregated into allele-specific summary statistics. These aggregate allelic fractions and standard deviations at each genomic position are stored in the Incremental Database (IncDB), which does not contain any patient-specific information. The 99.9% confidence interval for expected standard deviation at each allelic fraction was generated. Genomic positions where the observed standard deviation was below the confidence interval expected were cataloged as “suspect loci”, and mapped to variant calls in clinically relevant genes. Prioritisation of genes for diagnostic and reporting purposes can be adjusted according to the presence of suspect loci.

The observed relationship between standard deviation and the aggregate allelic fraction at each genomic locus was compared to the expected distribution assuming inherited variants in Hardy-Weinberg equilibrium in **Fig. 2A-C**. We labelled the positions which fell below the 99.9% confidence interval of the Mendelian model as ‘suspect loci’; ∼1-3% of all autosomal loci were suspect loci for at least one allele, which we define as unique suspect loci in this study. The upper and lower 99.9% confidence interval of the expected Mendelian model are illustrated in **Supplemental Fig. S1**. In all five data sets, suspect loci are mostly low allelic fraction (up to ∼40%), but with much lower standard deviations across samples compared to Mendelian variants with the same allelic fractions. The distribution of allelic fractions at suspect loci are illustrated in **Supplemental Fig. S2** for data sets 1-3. Unique suspect loci were counted across data sets 1 and 2, and the overlap revealed that most unique suspect loci were not shared between both pipelines (**Fig. 2D**). Single nucleotide variants (SNVs) that were called in the gold-standard reference NA12878, an individual sample separate from all of the data sets used, were analysed to check if they validated both suspect locus positions and their corresponding suspect alleles in data sets 1 and 2 (**Fig. 2E**). There were 147,000 SNVs reported in NA12878 that were also annotated as suspect in both data sets 1 and 2, which corresponded to an overlap proportion roughly twice as high as **Fig. 2D**, showing that suspect variant calls were more frequent at suspect loci present in both data sets than in either one data set alone.

**Figure 2:**
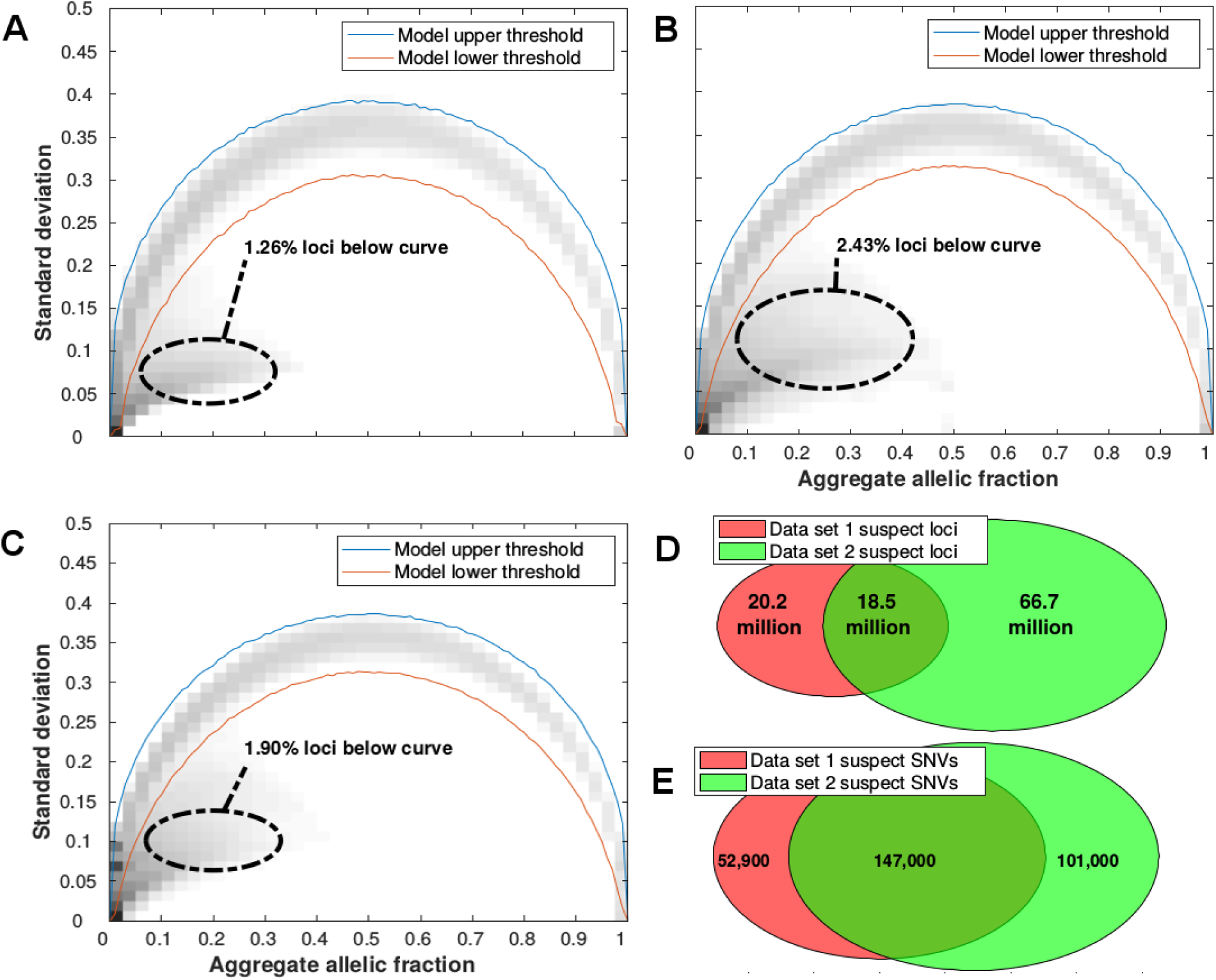
Identification of suspect autosomal loci/allele combinations with persistent low allelic fractions across patients. Observed and expected alternative allele fractions were estimated from five different whole genome sequenced cohorts: (**A**) Data set 1, (**B**) Data set 2, (**C**) Data set 3. For all loci in autosomal chromosomes, the standard deviation and aggregate allelic fraction values from the IncDB were plotted against each other in a density plot. The darker regions have the highest concentration of loci. The red lines indicate the upper and lower boundaries of the 99.9% confidence interval for the expected allelic standard deviation (shown in Supplemental Fig. S1A,B respectively). Suspect loci for each cohort were defined as the loci with standard deviation below their simulated model lower threshold. A total of 2.8 billion autosomal loci were assessed. (**D**) Venn diagram showing the overlap of all suspect loci between data sets 1 and 2. (**E**) Venn diagram of overlap at suspect loci where SNVs have been called. A total of 3.44 million autosomal SNVs were not annotated as suspect in either data set 1 or 2.

### Enrichment of suspect loci within specific genomic regions

Unique suspect loci were found to be present across the entire sequenced autosomal genome and only absent in unsequenced sections, although their prevalence varied across sequenced regions both locally and showing larger trends across chromosomes (**Fig. 3**). We examined the distribution of unique suspect loci across different regions of the genome (**Fig. 4**), and recorded the regional enrichment of unique suspect loci (**Supplemental Table S1**) using odds ratios (OR). All odds ratios calculated were highly statistically significant due to the very large number of genomic positions sampled, even when the odds ratios were close to 1. The 95% confidence interval lower and upper bounds were both equal to the reported odds ratios to 3 significant figures in all cases. The highest/least significant p-value recorded was for the enrichment of suspect loci in the intellectual disability gene panel in data set 2 (OR=1.01, p=8.69×10^−41^). All other p-values ranged from 10^−322^ to 10^−79^.

**Figure 3:**
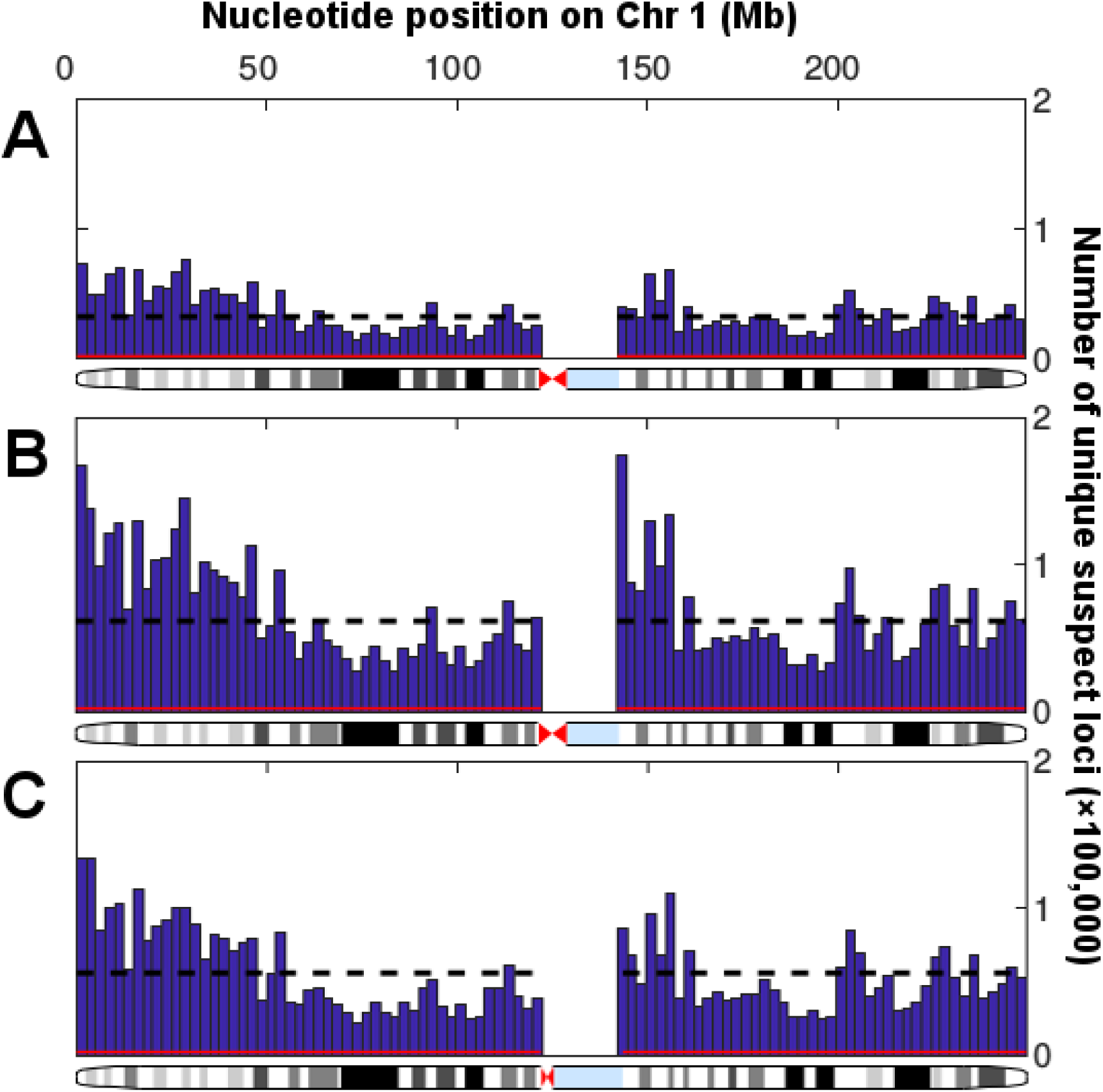
Variability in distribution of unique suspect loci in sequenced regions of Chromosome 1. Histograms show the numbers of suspect loci in 100 regular intervals across Chromosome 1, with the number of suspect loci per 2.49 million bp bin on the y axis and the nucleotide position on the x axis. There were no suspect loci at the centromere since this could not be sequenced. The black dotted line shows the mean number of suspect loci per bin, while the red line shows the number of suspect loci in each bin that would be expected by chance (1 per 1000 loci). (**A**) Data set 1, Personalis IncDB and GRCh37. (**B)** Data set 2, 100,000 Genomes Project and GRCh37. (**C**) Data set 3, 100,000 Genomes Project and GRCh38.

**Figure 4:**
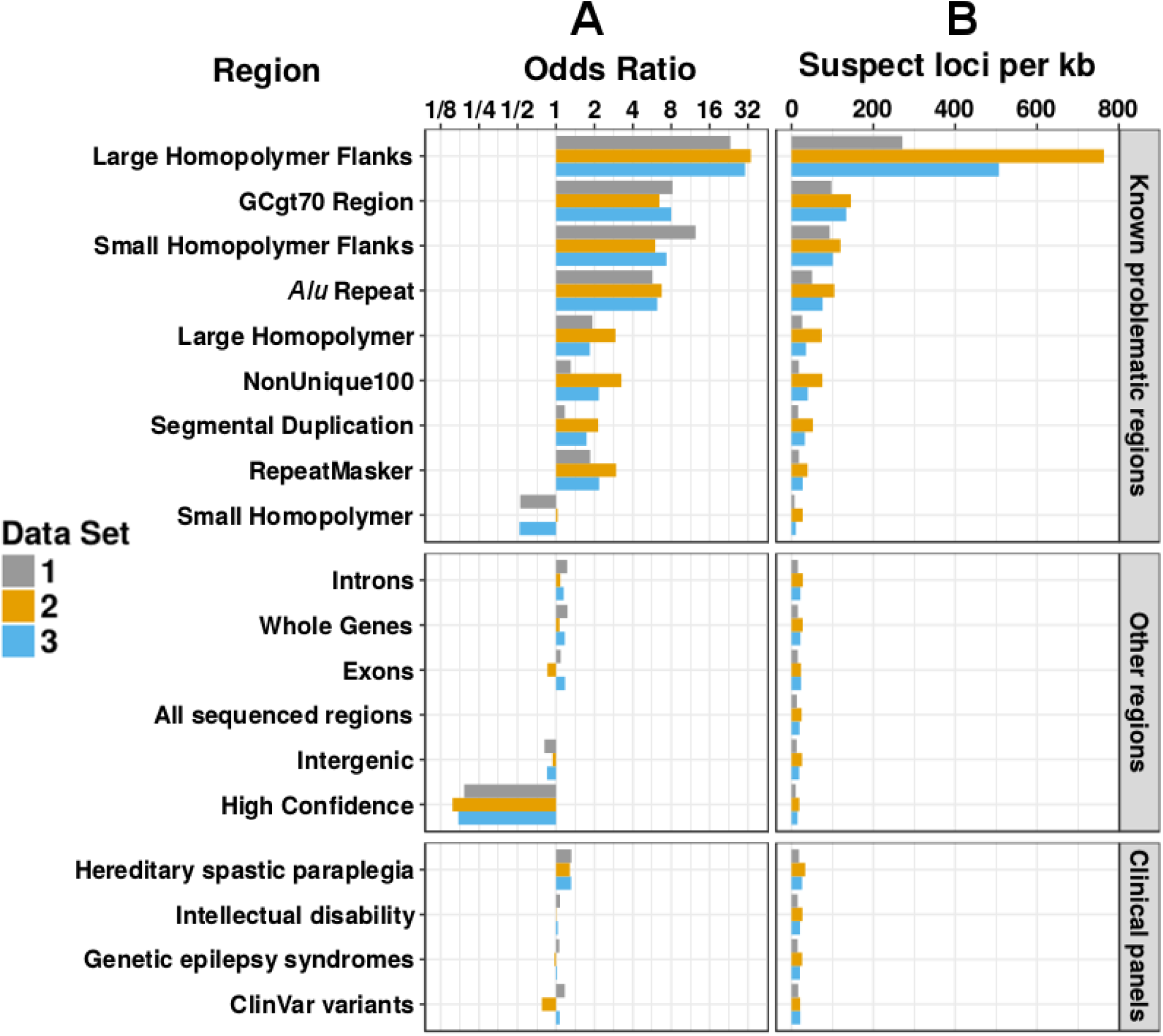
Enrichment of unique suspect loci in different types of genomic regions. All odds ratios shown are statistically significant and equivalent to the 95% upper and lower confidence intervals to 3 significant figures. (**A**) *Log*_2_ scale barplot showing the odds ratios for regional enrichment of unique suspect loci across all autosomal chromosomes from three different data sets. All odds ratios were calculated and shown to be significant using Fisher’s exact test. Regions were compared to “All sequenced regions” (odds ratio of 1). (**B**) Barplot showing the number of unique suspect loci per kb across all autosomal chromosomes from three different data sets. Although clinical and high confidence regions had a lower rate of suspect loci per kb, over the entire genome they still contained a large number of suspect loci overall in high confidence regions (24.1/45.1/34.6 million in data sets 1, 2 and 3) and clinical regions (2.55/4.64/3.55 million in data sets 1, 2 and 3 within the intellectual disability panel alone).

The 100 base pair flanks of large (≥ 20 bp) homopolymers were the most heavily enriched for suspect loci by a large amount (OR = 23.29, 33.69, 30.37 for data sets 1, 2, 3). Small (<20 bp) homopolymers’ 100 base pair flanks (OR = 12.40, 5.99, 7.36), GC-rich regions (OR = 8.19, 6.47, 7.99) and *Alu* repeats (OR = 5.70, 6.74, 6.21) were also strongly enriched for suspect loci. Large homopolymers (OR = 1.93, 2.92, 1.84) and the RepeatMasker regions (OR = 1.85, 2.96, 2.18) were mildly enriched for suspect loci in data sets 1-3. Small homopolymers on the other hand were depleted or unenriched (OR = 0.525, 1.03, 0.519), and the NIST GIAB high confidence region was strongly depleted (OR = 0.191, 0.154, 0.172) for suspect loci. The NonUnique100 region showed much greater enrichment of suspect loci in data sets 2 (OR = 3.26) and 3 (OR = 2.17), than in data set 1 (OR = 1.30), and segmental duplications also displayed this (OR = 1.17, 2.15, 1.74).

### Systematic biases confirmed in the gold-standard reference sample

Our analysis based on aggregate allele fraction statistics found the presence of suspect loci in all cohorts. In order to confirm that these loci occur independently from the samples examined, specific suspect loci were examined in the reference sample NA12878 cell line, which was sequenced and had variants called by Personalis Inc. using the same pipeline as data set 1 for comparison, but which was not part of any of the data sets used to generate the IncDBs. We divided the SNVs called using this pipeline into groups based on whether they were annotated as suspect or non-suspect, whether they occurred in the NIST GIAB benchmark regions and whether they matched the v3.3.2 NIST GIAB benchmark variants at those positions.

Suspect SNVs, which accounted for ∼5% of all SNV calls, mostly reported low allele fractions (0.300 median compared to 0.579 in non-suspect SNVs) in the read pileup of NA12878, with the exception of suspect SNVs within the benchmark region that matched the benchmark variants. These were the least common type of suspect SNV and most closely resembled the allelic fraction distribution of non-suspect SNVs (1,358 SNVs, **Fig. 5A**), suggesting that suspect loci that matched the benchmark variants were likely to be false positives. Suspect SNVs within the benchmark regions usually did not match the benchmark variants within these high confidence regions (1,839 SNVs, **Fig. 5B**). Most suspect SNVs occurred outside of the benchmark regions (23,802 SNVs, **Fig. 5C**).

**Figure 5:**
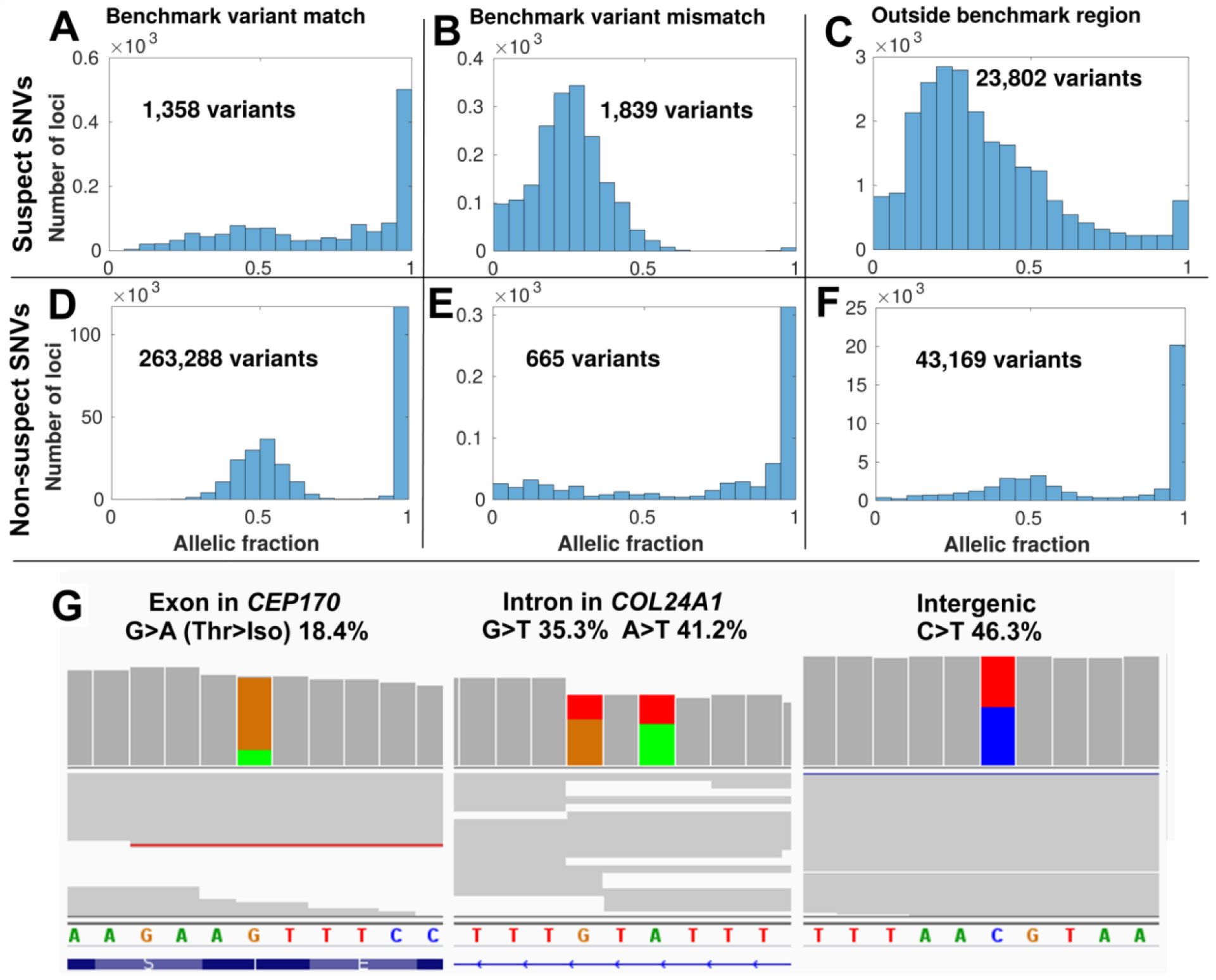
Suspect loci in detected variants of a gold-standard genome. Distribution of allelic fractions of SNVs called in Chromosome 1 of NA12878, classified as either suspect SNVs (top row - **A-C**), if they corresponded to suspect alleles in data set 1 (Personalis), or non-suspect SNVs (second row - **D-F**). SNVs were also classified based on whether they matched the NIST GIAB v3.3.2 benchmark variants (left column), didn’t match the benchmark variants (middle column) or were outside of the GIAB benchmark region (right column). Low coverage variants (<10 supporting reads) were excluded from this analysis. (**G**) Cropped panels from the Integrative Genomics Viewer (Robinson et al. 2017), highlighting suspect loci from data set 1 in Chromosome 1 which were called as variants separately in NA12878. NA12878 was sequenced with Illumina HiSeq, but not used as part of the patient data set to create the IncDB (Zook et al. 2014, 2019). Reads are shown in grey with coloured bands where non-reference allelic reads were observed (A=Green, C=Blue, G=Brown, T=Red). Suspect SNVs and their respective read proportions in the NA12878 cell line are indicated above - these systematically occur at similar levels across all patients in the IncDBs used to identify them. Left/Middle: Suspect SNVs in exonic and intronic regions of genes in the PanelApp Intellectual Disability panel (Martin et al. 2019). Right: Suspect SNV in an intergenic region.

In contrast, most called SNVs in NA12878 were non-suspect, were within the GIAB high confidence benchmark region and matched the GIAB benchmark variants for those positions (263,288 SNVs, **Fig. 5D**). These conformed to the expected distribution of allele fractions for heterozygous or homozygous SNVs, with peaks around 50% and 100%. Non-suspect SNVs that were inside the high confidence benchmark region, but which did not match the benchmark variants were the least common and only showed a peak around 100%, with no peak observed for heterozygous variants and low levels of SNVs at all allelic fractions, suggesting that these were sequenced the worst of the non-suspect loci (665 SNVs, **Fig. 5E**). Non-suspect SNVs outside of the benchmark regions were also numerous and had a similar pattern to non-suspect SNVs which matched benchmark variants, although the peaks were broader (43,169 SNVs, **Fig. 5F**), suggesting that the allelic fractions recorded for these were slightly less accurate.

Suspect loci called as SNVs occurred across all types of genomic region, including clinically-important positions (**Fig. 5G**). All together, this suggests that the systematic biases at suspect loci can contribute to false positive variant calls in clinically important regions, even within the NIST GIAB high confidence regions. However, suspect SNVs that match the v3.3.2 NIST GIAB benchmark variants should be treated with caution, since their allelic fraction distributions suggest that these may indeed be true variants, so the NIST GIAB benchmark variants are a necessary and complementary resource to combine with IncDB-based methods for annotating systematic error accurately.

## Discussion

The main aim of this study has been to develop and evaluate a novel statistical method to identify positions of the genome that are prone to systematic bias in genomic sequencing and alignment, using anonymised summary patient data. We developed an approach to quality control sequenced reads in the autosomal genome by cataloguing genomic positions that persistently present a low-fraction alternate allele across patients rather than reflecting true biological variation, which we have labelled as ‘suspect loci’. We have explored the extent to which these systematic biases occur across varied genomic regions, including regions known to be difficult to sequence, higher confidence regions and clinical panels. We have also confirmed the existence of these systematic biases in an independent gold-standard reference genome and the utility of our approach.

For the purposes of this paper, we have defined systematic biases as locus-specific errors in the accurate quantification of allelic fraction that affect all samples sequenced with the same pipeline to a similar extent. Within this definition, our method should successfully identify almost all single-nucleotide allelic fraction biases greater than 1.5%, but does not have the sensitivity to detect smaller biases at the confidence level chosen (**Supplemental Figure S2**). For example, using a confidence interval of 99.9% and a sample size of 150 (reflecting the parameters used to classify suspect loci for data set 1), ∼9.6% of sites with a 1% systematic error rate were classified as suspect loci, while ∼99.2/99.8/100/100% of sites with a 2/3/4/5% systematic error rate were classified as suspect loci respectively, when we simulated sites with these systematic error rates, representing a very high resolution. We have not tested this method for identifying systematic biases in the detection of indels or structural variants, but with modifications a similar approach could in theory be used to identify those types of systematic error. We carried out Monte Carlo simulations to check if this method could be used to identify CNVs, but found that CNVs were not annotated by this method below a population frequency of 10%, and could only be detected if intervening SNV alleles had unbalanced aggregate allelic fractions >10%. We expect that CNVs that fulfill these criteria are exceedingly rare so this is unlikely to cause much of the systematic bias this method identifies. Our method cannot be used for identifying types of systematic bias that are not locus-specific, such as general biases in the detection of certain alleles (e.g. GC bias (Chen et al. 2013)), biases that systematically affect the ends of genomic reads being sequenced (Ma et al. 2019), or other context-specific biases that do not occur in a systematic or locus-specific manner. For example, we would not expect to identify mosaicism unless this was systematic, with a high proportion of individuals having the same alternative genotypes at the same genomic coordinates in the same fraction of cells in their tissue. To evaluate whether age-related mosaicism could have been present in our results we compared the numbers of suspect loci (54.6 million) and suspect SNVs (43,832) in data set 5, which used a more elderly cohort of patients, with data set 3 (63.2 million suspect loci and 46,287 suspect SNVs) and data set 4 (90.3 million suspect loci and 59,634 suspect SNVs), which both used the same sequencing pipeline, but did not observe an increase in suspect loci in the gene *TP53*, which is thought to be correlated with this (Yizhak et al. 2019), or in general across the genome (**Supplemental Fig. S3**), so we could not find evidence for mosaicism affecting suspect locus annotation. The main aim of our approach is not to identify a specific cause of systematic bias, but rather to act as a quality control method, to catalogue where these systematic biases occur so that they can be filtered out of scientific and clinical results where they may lead to inaccurate conclusions. As a result, our study does not seek to identify why the systematic biases we have identified occur, even if some of our results suggest possible causes that could be followed up by future studies.

To investigate the extent to which differences in mutations between large patient cohorts affected the annotation of suspect loci when technical procedures were held constant, we compared the overlap between suspect loci for data sets 3-5, for which all patients were sequenced using the same protocol and pipeline. These 3 data sets showed considerable suspect loci overlap (mean overlap=65.7%) and suspect SNV overlap (mean overlap=71.9%) in the intersection of all 3 data sets, although there were some differences likely due to private mutations (**Supplemental Fig. S3**). Compared with the overlap between data sets 1 and 2, for which separate technical procedures were used (mean overlap=34.8% for suspect loci, 66.4% for suspect SNVs), the overlap was ∼30% greater for suspect loci, but only ∼5% greater for suspect SNVs, suggesting that differences in the technical procedures used played a large role in suspect locus annotation in general, but genetic differences between cohorts were particularly important when identifying systematic biases that could affect variant calls. We would therefore recommend users increase the size of IncDBs they generate as much as is reasonably possible, to maximise the numbers of systematic biases they can detect by this method, at least until overlap between separate IncDBs approaches a percentage.

For loci unaffected by systematic bias, the standard deviation and aggregate allelic fraction stored in the IncDB were expected to relate to each other in accordance with Hardy-Weinberg equilibrium. However, ∼1-3% of autosomal loci had significantly lower standard deviation than predicted by Hardy-Weinberg equilibrium (p=0.0005), suggesting the presence of systematic bias across numerous genomic loci. At these loci most individuals appear to present a low-fraction allele. Persistent low-fraction alleles would be inconsistent with our understanding of human genetics and are presumably a technological artifact: a bias or systematic error in the sequencing technology itself, or perhaps in the read mapping. The impact of these suspect loci is magnified in the context of studies looking at large numbers of genomic positions, since a small percentage of this would still correspond to a high number of genomic positions affected by systematic bias. It is therefore clear that these systematic biases are of concern and deserve further attention. Previous studies examining systematic sequencing bias have primarily focussed on biases in total coverage across loci and have not examined position-dependent systematic biases in the allelic fraction (Ross et al. 2013; Cheung et al. 2011), therefore, no previous estimates for the prevalence of systematic sequencing bias in allelic fractions were available to compare with our estimate of ∼1-3% of all autosomal loci in the human genome.

Our estimate indicated a very large number of genomic sites affected by systematic bias, potentially more than might be expected, so we analysed what proportion of these would be filtered out by applying a strongly conservative confidence threshold (99.999%) in addition to the default threshold used in this study (99.9%) to data sets 1-3 and compared how this affected suspect locus numbers, allelic fractions and overlap between data sets 1 and 2. This filtered out a very high proportion (99.98/91.06/92.68% in data sets 1, 2 and 3 respectively) of the suspect loci that had allelic fractions <0.03, since it was rare for loci exhibiting very small systematic biases to have a standard deviation lower than the 99.999% confidence threshold (**Supplemental Fig. S4**) compared to the 99.9% confidence threshold (**Supplemental Fig. S2**). In contrast, a much lower proportion (25.95/17.74/23.40% in data sets 1, 2 and 3 respectively) of the suspect loci with larger systematic biases (allelic fractions>0.03) were filtered out at the 99.999% confidence threshold. Since it is the larger systematic biases that are most likely to affect variant calling, this shows that the method is robust for identifying most important systematic biases, even at a strongly conservative confidence level. The percentage overlap of suspect loci between data sets 1 and 2 decreases slightly when a more conservative threshold (99.999% confidence interval) is used for defining suspect loci (from 47.9/21.7% to 41.1/16.1% of data set 1 and 2 respectively, **Supplemental Fig. S5**).

Suspect loci were widespread across all sequenced chromosomal segments, but there was variability between different types of genomic region. Despite this, there was little or no depletion of suspect loci in some regions expected to have more accurate sequencing, such as exons and the clinical gene panels, suggesting that greater caution in these areas is justified when calling variants. The NIST high confidence regions displayed the greatest depletion of suspect loci as expected and had the lowest proportion of suspect loci per kb, while homopolymer flanks displayed the greatest enrichment of suspect loci.

The correlation between chromosomal suspect locus density and the proportions of chromosomes within large homopolymer flanks was high (Spearman’s Rho = 0.955, 0.861, 0.820, p = 5.11×10^−12^, 2.76×10^−7^, 3.07×10^−6^ for data sets 1, 2 and 3) respectively, accounting for the majority of the variability in the proportion of suspect loci in autosomal chromosomes (**Supplemental Fig. S6**). The flanks of large homopolymers are particularly prone to errors in alignment compared to other types of sequencing error, especially for Illumina sequencing (Laehnemann et al. 2016) as was done here, suggesting that the misalignment of reads could be a source of systematic biases.

100bp was chosen as the flank length because it is of the same order of size as the read lengths used for the Illumina sequencing and we would not expect anything outside of 100bp from a homopolymer to be covered by sequencing reads, so the 100bp length is the theoretical maximum length at which we might expect effects to occur. Homopolymers cannot affect the sequencing beyond 1 read length from their edge. In practice most suspect locus enrichment occurred within the 3bp flanking regions however, with enrichment odds ratios greatly increasing when the flank size was reduced to 3bp in both large homopolymers (from 23.3/33.7/30.4 in data sets 1, 2 and 3 respectively in 100bp flanks, to 171.3/94.3/97.6 in 3bp flanks) and small homopolymers (from 12.4/6.0/7.4 in 100bp flanks in data sets 1, 2 and 3 respectively, to 30.9/12.2/13.3 in 3bp flanks).

In addition we found differences in regional systematic biases between data sets. For example, the NonUnique100 region showed much greater enrichment of suspect loci in data sets 2 (OR = 3.26) and 3 (OR = 2.17), than in data set 1 (OR = 1.30). This region is defined as containing all sequenced positions where a single 100-bp read cannot map uniquely, so we would expect a large proportion of systematic biases here to mainly correspond to alignment errors rather than sequencing chemistry. The higher enrichment of suspect loci in NonUnique100 in data sets 2 and 3 could therefore indicate that the different aligners used (Isaac aligner (Raczy et al. 2013) for data set 2 and 3 rather than BWA-MEM (Li 2013) for data set 1) could be the main explanation for the different levels of systematic bias found between the data sets in this region. The Isaac aligner is known to be a faster aligner than BWA-MEM, but previous validation attempts found that it was slightly worse than BWA-MEM in terms of accuracy, especially outside of NA12878 (Mainzer et al. 2015). The increased frequency of systematic biases we detected with the Isaac aligner in data sets 2 and 3 therefore seems to indicate that BWA-MEM is still superior in terms of accuracy, at least compared with the versions of the Isaac aligner used (SAAC00776.15.01.27 and iSAAC-03.16.02.19 respectively).

We also found great variability in the distribution of suspect loci within individual genes tested (**Supplemental Data S1, S2**). For example, within the clinical gene panels there were very low proportions of suspect loci (0% out of 2,021 total loci in *LIPT2* in data set 2) to very high proportions of suspect loci (36.6% out of 3,759 total loci in *NPRL2* in data set 2), suggesting that some genes might be particularly prone to systematic sequencing biases. This is likely because the genes intersect with the problematic regions we examined in different ways. Both of these genes are associated with genetic epilepsy syndromes, but *LIPT2* is not affected by systematic sequencing biases at all, while these heavily affect *NPRL2*. Clinicians focussing on specific genes for diagnostic purposes could use this information to identify how much caution they need when assessing pathogenic variants in those genes (**Table 2**). These suspect loci were confirmed in an independent reference sample sequenced using the same pipeline, including within introns, exons, intergenic regions and NIST GIAB high confidence regions, and within called variants in clinically relevant regions such as the PanelApp disease panels (Martin et al. 2019). In addition, we demonstrated that our approach could be used in combination with the NIST GIAB benchmarking variants to improve suspect locus annotation by identifying which suspect SNVs were likely to be false positives.

**Table 2:** Verified pathogenic variants in ClinVar that are included in diagnostic gene panels and at locations of suspect loci found in data sets 3-5.

| dbSNP ID | Gene Affected | Variant Type | Diagnostic Panel | ClinVar Associated Disease |
| --- | --- | --- | --- | --- |
| rs1131691563 | *AMPD2* | splice donor variant | Ataxia and cerebellar anomalies | Not provided |
| rs1553201258 | *DARS2* | intron variant | Ataxia and cerebellar anomalies | Cerebral cortical atrophy |
| rs370132645 | *OTOF* | splice donor variant | Auditory Neuropathy Spectrum Disorder | Rare genetic deafness |
| rs587779190 | *MSH2* | stop gained | Adult solid tumours cancer susceptibility | Lynch syndrome |
| rs63750640 | *MSH2* | missense variant | Adult solid tumours cancer susceptibility | Lynch syndrome |
| rs1114167845 | *MSH2* | stop gained | Adult solid tumours cancer susceptibility | Lynch syndrome |
| rs587779197 | *MSH2* | missense variant | Adult solid tumours cancer susceptibility | Lynch syndrome |
| rs587779195 | *MSH2* | splice donor variant | Adult solid tumours cancer susceptibility | Lynch syndrome |
| rs121918737 | *SCN1A* | missense variant | Genetic epilepsy syndromes | Severe myoclonic epilepsy in infancy |
| rs267607819 | *MLH1* | splice acceptor variant | Adult solid tumours cancer susceptibility | Lynch syndrome |
| rs1559551570 | *MLH1* | frameshift | Adult solid tumours cancer susceptibility | Hereditary nonpolyposis colon cancer |
| rs148891849 | *DNAH5* | stop gained | Primary ciliary disorders | Primary ciliary dyskinesia |
| rs7755898 | *CYP21A2* | stop gained | Disorders of sex development | Classic congenital adrenal hyperplasia |
| rs1554247637 | *ARID1B* | frameshift | Coffin-Siris syndrome | Inborn genetic diseases |
| rs1562671039 | *PMS2* | stop gained | Adult solid tumours cancer susceptibility | Not provided |
| rs193929376 | *GCK* | splice donor variant | Diabetes - neonatal onset | Permanent neonatal diabetes mellitus |
| rs121909112 | *HSPB1* | missense variant | Distal myopathies | Charcot-Marie-Tooth disease type 2F |
| rs111033565 | *PRSS1* | missense variant | Pancreatitis | Hereditary pancreatitis |
| rs794728419 | *KCNH2* | splice donor variant | Short QT syndrome | Not provided |
| rs587777641 | *GPIHBP1* | missense variant | Severe hypertriglyceridaemia | Hyperlipoproteinemia, type ID |
| rs1563963464 | *APTX* | splice acceptor variant | Ataxia and cerebellar anomalies | Ataxia-oculomotor apraxia type 1 |
| rs146292819 | *ABCA1* | missense variant | Hereditary neuropathy | ABCA1-Related Disorders |
| rs864321692 | *WAC* | stop gained | Intellectual disability | Desanto-shinawi syndrome |
| rs587782455 | *PTEN* | splice acceptor variant | Adult solid tumours for rare disease | PTEN hamartoma tumour syndrome |

We further confirmed that suspect variants were filtered out of the Genome Aggregation Database (gnomAD) using their quality control processes (**Supplemental Fig. S7**). The gnomAD database is a large database of all of the variation found across a large ethnically diverse population, taken from 125,748 exomes and 15,708 genomes (Karczewski et al. 2019). Our results revealed that suspect variants were also widespread in the gnomAD database, even after filtering by gnomAD’s quality control process, across allelic frequencies. We also evaluated whether suspect loci could be identified simply by using a quality threshold (**Supplemental Fig. S8**). Non-suspect loci had significantly higher proportions of high quality reads, with >90% of reads having sequencing and mapping quality scores >20 in ∼90% of non-suspect loci. This demonstrated that low quality reads were more frequent among suspect loci to a large degree, suggesting that improving sequencing and alignment quality could help with decreasing these systematic biases. However, there was still significant overlap between read quality at suspect and non-suspect loci. Finally, we analysed whether read depth could be used to quality control for suspect loci (**Supplemental Fig. S9**). Kolmogorov-Smirnov tests confirmed that there was a different distribution of read depths between the sets of genomic positions with and without suspect loci (D=0.4289, 0.2885, 0.3079 and p<10^−15^ for data sets 1,2,3 respectively), but there wasn’t a clear correlation between read depth and likelihood of a position having suspect loci. While the lower quartile and median values for read depth were slightly lower for unique suspect loci, for data sets 2 and 3 the upper quartiles were significantly higher at unique suspect loci. We could only conclude that there was a greater breadth of read depth in positions annotated as unique suspect loci than other positions. A possible explanation for our results could be that most unique suspect loci have slightly lower read depths than non-suspect loci, but some unique suspect loci have extremely high read depth, perhaps as a result of systematic alignment errors attributing reads from multiple locations in the genome to the same genomic position at these suspect loci. Our analyses therefore suggest that the existing read depth thresholds and quality control procedures commonly used in sequencing would not be sufficient to filter out the systematic biases, and the reported variants in large population studies such as gnomAD may need to be reassessed.

We have demonstrated the utility of IncDBs to assess the quality of clinical whole genomes of five independent cohorts sequenced by commercial and public healthcare organisations while maintaining patient anonymity. In addition to showing the utility of this approach on whole genome Illumina sequencing, IncDBs could be applied to data from different types of sequencing platforms in the future, including specific targeted, exome-sequencing and long-read technologies such as PacBio and Oxford Nanopore. Our agile approach for detecting suspect loci could be deployed in various settings where the raw data for individual genomes cannot be accessed, for instance when patient confidentiality must be maintained. Under those conditions, being able to identify systematic biases would enable improvements to variant calling and has the potential to reduce errors in clinical genomic testing.

## Methods

### Data sources

#### Data set 1 (Personalis Inc.)

WGS data were obtained by Personalis Inc. using Illumina HiSeq the standard library prep and sequencing protocol. Paired-end reads of 100bp length was mapped with BWA-MEM (Li 2013) to align reads against the GRCh37 reference human genome (**Table 1**). The mean depth of coverage across patients was 45×. There were 150 non-cancer individuals in the cohort, including triplets (infant and two parents) recruited from hospitals in the USA and a mix of ethnicities, but specific age or ethnic data were not available.

#### Data set 2 (100,000 Genomes Project)

Blood samples were taken from 215 distinct patients of mixed ethnicities with non-cancer neurological diseases in each cohort, recruited from hospitals in the UK. The libraries were prepared using Illumina TrueSeq DNA PCR-Free Library Prep for the majority of the samples. For a small proportion of samples, when the concentration was less than 20ng/µl at Illumina, the Nano DNA library prep method was used. All main programme samples were sequenced using 150bp length paired end sequencing on HiSeqX and the mean depth of coverage across patients was 30×. WGS data were obtained by Genomics England’s 100,000 Genomes Project, using Illumina HiSeq sequencing and mapped with the Illumina Isaac-aligner (version SAAC00776.15.01.27) (Raczy et al. 2013) to align reads against the GRCh37 reference human genome. All variants were called using Isaac Variant Caller (Starling, v2.1.4.2) (Raczy et al. 2013)and annotated using Ensembl database (v72) (Cunningham et al. 2019). The median age of patients was 13 (**Supplemental Fig. S10**).

#### Data sets 3, 4, 5 (100,000 Genomes Project)

No patients were in multiple data sets. Blood samples were collected and library prepped in the same way as data set 2. WGS data were obtained by Genomics England’s 100,000 Genomes Project, using Illumina HiSeq sequencing and mapped with the Illumina Isaac-aligner (iSAAC-03.16.02.19) to align reads against the GRCh38 reference human genome. The reads were aligned by Illumina with Issac (v03.15.09.04) and variants called by Isaac Variant Caller (Starling, v2.3.13) (Raczy et al. 2013) with annotation by Ensembl (v81) (Cunningham et al. 2019). Data sets 3, 4 and 5 all used 215 distinct patient samples from the non-cancer neurological diseases cohort, but the patients in data sets 3 and 4 were both randomly sampled so these data sets can be treated as technical replicates. For data set 5 we used data from the same cohort, but selected all of the oldest patients available since the patients in data sets 3 and 4 were generally very young. The age distributions of patients in cohorts 2-5 are illustrated in **Supplemental Fig. S10**, and had median ages of 13, 16, 13 and 64 respectively. The numbers of patients across different ethnic categories in data sets 2-5 were recorded in **Supplemental Table S2**.

### Incremental Database Generation

The read depth values for each allele (A, C, G, T) at every autosomal genomic locus were calculated from aligned BAMs and divided by the total read depth at the corresponding loci to get the allelic coverage fraction, *x*_*p*_, for each allele at each locus in each patient. Individual IncDBs were created for each data set from the aggregate allelic fraction and standard deviation values for each allele at each locus across the entire cohort, which were calculated from *x*_*p*_ as described below:

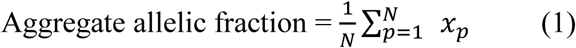

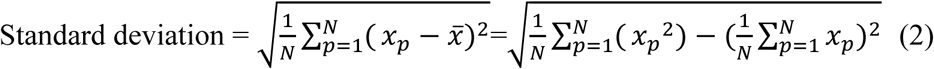

*N* is the number of patients, p is the patient identifier, *x*_*p*_ is the allelic coverage fraction for a specific allele in patient p, and 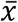 is the mean of all of the allelic coverage fractions for that same allele across all patients (aggregate allelic fraction).

Notice that to compute the aggregate allelic fraction, we do not store each individual’s *x*_*p*_ values, but the sum of *x*_*p*_ across all individuals. Similarly, we can compute the standard deviation across individuals by storing the sum of *x*_*p*_, as well as the sum of *x*_*p*_ ^2^. This approach not only removes all individual-specific genomic information, but also allows the IncDB to grow indefinitely, as more samples are sequenced and analyzed: they can simply contribute to the running sums of *x*_*p*_ and *x*_*p*_ ^2^. Also note that the sums in the equations above do not take up more size on disk as the number of samples increases, so the overall IncDB file size does not increase as new samples are added.

### Identifying loci affected by systematic bias (suspect loci)

For each locus in all autosomal chromosomes, the standard deviation and aggregate allelic fraction values were taken from the IncDB and plotted against each other in a density plot using MATLAB 9.6 (R2019a). The main bow-shaped feature of this plot (**Fig. 2A-C**) was the expected result of Mendelian alleles in the human population, present at a variety of population frequencies, while observed loci with standard deviations below the 99.9% expected confidence interval for a given nucleotide were defined as suspect loci for that allele. Autosomal positions that displayed at least one suspect allele at that position were termed unique suspect loci. The total count of unique suspect loci was therefore lower than the total count of allele-specific suspect loci, since some positions had multiple suspect alleles.

The 99.9% confidence interval was estimated using Monte Carlo sampling as detailed in the pseudocode below. Monte Carlo sampling used 3 nested loops which respectively simulated the standard deviation at a single genomic locus between n individual patient allelic fractions (loop 3), 1000 times to calculate the upper and lower 99.9% confidence intervals (loop 2), for each aggregate allelic fraction from 0 to 1 in intervals of 0.01 (loop 1). The standard deviation values were recorded and used to classify suspect loci as visually illustrated (**Fig. 2A-C**). n=150 for data set 1, and n=215 for data sets 2/3. The model assumed an error rate of 0.01, corresponding to an approximation of the error rate of Illumina WGS (Wall et al. 2014). Approximately 90% of genomic reads in data set 1 had a quality score of 20 or above, corresponding to this error rate (**Supplemental Fig. S8**). Decreasing the assumed error rate increases the numbers of sites with very low systematic allelic biases that get annotated as suspect loci, by increasing the standard deviation threshold for classification as suspect (**Supplemental Fig. S11**). The distribution of the suspect locus allelic fractions at an assumed error rate of 0 are illustrated in **Supplemental Fig. S12**, for comparison with the allelic fractions at an assumed error rate of 0.01 (**Supplemental Fig. S2**).

**

Monte Carlo simulation of standard deviation (pseudocode)

~~~
1. for aggregate allelic fractions, AAF, from 0 to 1 in intervals of 0.01 (each representing a 
simulated single autosomal genomic position with that aggregate allelic fraction across all 
patients) do {
   2. repeat 1000 times {
       3. repeat for n simulated patients {
       Randomly generate diploid genotype for each simulated patient using the
       binomial distribution (assuming two alleles) at the given AAF value;
       Assuming a sequencing error rate of 0.01, randomly
       draw c reads from the binomial distribution to simulate observed major/minor allelic reads for the 
       simulated diploid genotype;
       Divide by total read depth, c, to get the individual allelic fractions for each 
       patient;
       }
   Calculate the standard deviations between the individual allelic fractions for all n patients at the simulated genomic position;
   }
Maximum and minimum values of 1000 repetitions mark upper and lower 99.9% confidence intervals for standard deviation at given AAF;
}
**
~~~

### Analysis of regional enrichment of unique suspect loci

Histograms of suspect locus density across Chromosome 1 were plotted in MATLAB alongside chromosome ideograms taken from the UCSC Genome Browser (Kent et al. 2002) (GRCh37 http://hgdownload.cse.ucsc.edu/goldenpath/hg19/database/cytoBandIdeo.txt.gz; and GRCh38 http://hgdownload.cse.ucsc.edu/goldenpath/hg38/database/cytoBandIdeo.txt.gz) in order to show suspect locus density in comparison with chromosomal banding patterns.

BED files for 18 different types of genomic region were analysed to check enrichment of unique suspect loci using a Fisher’s exact test to calculate the exact significance values. Full contingency tables for all regions in data sets 1-3 (autosomal chromosomes only) are available in **Supplemental Table S1** to show how these were calculated.

The regions tested were the NIST GIAB high confidence regions, *Alu* repeats, GCgt70 (> 70% GC content) regions, NonUnique100 regions (defined as all regions where a single 100bp read could not map uniquely), segmental duplications, small/large homopolymers, flanking regions of small/large homopolymers, the RepeatMasker region, introns, exons, genes, intergenic regions, ClinVar short variants and three neurological clinical panels (see next section for full list and details of BED files used).

“All sequenced regions” referred to all genomic loci where the number of aligned reads was greater than zero. There were no suspect loci outside of this region by definition, so the Odds Ratio was 1 by default.

### Genomic region BED file sources

NIST GIAB high confidence region (Xiao et al. 2014; Zook et al. 2014, 2019) - A selection of genomic loci covering the majority of the human genome that are considered to have high confidence calls (ftp://ftp-trace.ncbi.nlm.nih.gov/giab/ftp/release/NA12878_HG001/latest/GRCh37/HG001_GRCh37_GIAB_highconf_CG-IllFB-IllGATKHC-Ion-10X-SOLID_CHROM1-X_v.3.3.2_highconf_nosomaticdel.bed).

NonUnique100 - All regions where a single 100bp read cannot map uniquely (so all stretches on the reference that are 100bp or longer that are repeated on the GRCh37 reference).

Segmental duplication - Long DNA sequences (> 10kb) that are found in multiple locations across the human genome as a result of duplications.

GCgt70, small/large homopolymers and their 100bp flanking regions were all calculated as described below and BED files (available at https://doi.org/10.15131/shef.data.9927053) were generated in-house rather than downloaded from another source.

GCgt70 (GC content > 70%) - Regions with greater than 70% GC content. Loci were annotated as within this region if the surrounding 100bp around each locus had greater than 70% GC content.

Small/large homopolymer - Region of DNA containing a single nucleotide (9-19bp for small homopolymers, ≥ 20bp for large homopolymers).

Small/large homopolymer flanks - 100bp flanks surrounding the small/large homopolymer regions respectively.

RepeatMasker region - A BED file containing a variety of different types of repeats (Smit, AFA, Hubley, R & Green, P 2013-2015). The open-3-2-7 version of RepeatMasker was downloaded from the UCSC Table Browser (Karolchik et al. 2004) (https://genome.ucsc.edu/cgi-bin/hgTables).

*Alu* repeats (Hasler and Strub 2007) - The most common type of transposable element in the human genome, of which there are over one million copies. The BED file was composed of all RepeatMasker Regions downloaded from the UCSC Table Browser (https://genome.ucsc.edu/cgi-bin/hgTables) that were annotated as *Alu* repeats in the repName column.

BED files were also downloaded for genic regions, intergenic regions, exonic regions, intronic regions (03/22/19) and ClinVar short variants (06/12/19), acquired from the UCSC Table Browser (https://genome.ucsc.edu/cgi-bin/hgTables).

Clinical panel BED files were also downloaded for the three most reviewed neurological clinical panels on PanelApp (Martin et al. 2019) (for intellectual disability (10/19/18), genetic epilepsy syndromes (02/07/19) and hereditary spastic paraplegia respectively (02/07/19) https://panelapp.genomicsengland.co.uk/panels/).

### Calculating allelic fractions at suspect loci in NA12878

An indexed BAM file for NA12878 Chromosome 1, sequenced using the same pipeline used in data set 1, was provided by Personalis Inc. We used the Integrative Genomics Viewer (IGV) (Robinson et al. 2017) to examine NA12878’s read pileup at data set 1 suspect locus positions at which Chromosome 1 SNVs had previously been called, to confirm that NA12878 exhibited low-fraction alleles at these positions. NA12878 was not part of any cohorts used to build the IncDBs in this study. Chromosome 1 SNVs in NA12878 were extracted from a VCF file corresponding to the sequencing pipeline used for data set 1, which was also provided by Personalis Inc. SNVs were classified as suspect if they corresponded to the same alleles at the same positions as suspect loci calculated for data set 1. The allelic fractions for these variants were calculated from the NA12878 BAM file using SAMtools v1.9 (http://www.htslib.org/download/) mpileup (Li et al. 2009; Li 2011). Variants with fewer than 10 supporting reads were deemed to have insufficient coverage and were not included.

### Analysing the proportion of gnomAD SNVs that are suspect

A list of all gnomAD variants (Karczewski et al. 2019), along with their allelic fraction and annotation as PASS-flagged or not, was obtained from a TSV file (available at https://doi.org/10.15131/shef.data.9927062). This was filtered to only include autosomal SNVs. These were classified as suspect and non-suspect SNVs as above. Variants in gnomAD were annotated as PASS variants if they were marked this way in gnomAD v2.1 (https://macarthurlab.org/2018/10/17/gnomad-v2-1/).

### Comparing sequencing quality between suspect and non-suspect loci

The coverage of different allelic reads across all loci/nucleotide combinations on Chromosome 1 was available for data set 1 (Cov), along with the corresponding coverage of allelic reads filtered to only include reads with sequencing and mapping quality scores greater than 20 (Cov20). The filtered coverage values of allelic reads (Cov20) were divided by the corresponding unfiltered coverage values (Cov) to get the proportion of allelic reads with sequencing and mapping quality scores both greater than 20 at each locus/nucleotide combination. The cumulative distributions of these values were calculated separately for locus/nucleotide combinations that were annotated as suspect loci or non-suspect loci.

### Using suspect loci to check the quality of your own sequenced samples

To address the limitations of existing quality control procedures used in WGS and variant calling pipelines discussed in this study, we have created resources for researchers and clinicians to carry out quality control of suspect variants occurring at positions that show a consistent systematic allelic bias (see Data Access). We have provided BED files containing all of the suspect loci that we have identified in all five data sets used for every allele, along with their corresponding aggregate allelic fractions, standard deviation between individuals’ allelic fractions, and allelic fractions corresponding to z values of −2, −1, +1 and +2. We suggest that researchers and clinicians using sequencing pipelines that are similar to any of the three pipelines used in this study, identify the corresponding BED files and check the allelic fractions of any variants that they have previously called that are present in these BED files. If the allelic fraction of a called variant is not significantly greater than would be obtained by systematic bias alone, i.e. if it is lower than the suspect allelic fraction at a z value of +1 for a milder filter or +2 for a stricter filter, then we would advise annotating or even removing that called variant. Researchers can also design custom filters based on their own preference if they wish to use different z value thresholds or other combinations of information. Researchers who are using Illumina WGS pipelines that have less in common with the pipelines shown here may also use a more conservative BED file of suspect loci to filter their data, containing only suspect loci found in both data sets 1 and 2, which is also included in the data set.

The suspect loci BED files provided by this study were only generated for specific WGS pipelines using Illumina HiSeq, and it is unlikely that these results could be used reliably for quality control of sequencing and alignment pipelines that are not similar to these. We would therefore recommend that researchers who have access to sequence data from a large cohort of patients should develop their own IncDB and calculate suspect loci for their sequencing pipelines based on that. The drawback of this, as opposed to using the suspect loci BED files above, is that this is computationally demanding and requires that researchers have access to large sequence data sets for their pipeline.

## Data access

Links to the sources of all non-confidential data used in this manuscript are referenced in the text at first mention and also below. We do not provide public links to download the raw sequence data we used in this study to generate the IncDBs for individual genome sequences since these are confidential. Data set 1 individuals were sequenced for commercial purposes. While raw sequence reads are no longer accessible for data set 1, we provide summary statistics from the BAM files stored in the Incremental Database (link below). In order to protect participants’ in data set 2 and 3, the 100,000 Genomes Project data can only be accessed through a secure Research Environment. To access the data for research, you must be a member of the Genomics England Clinical Interpretation Partnership (GeCIP), the research community set up to analyse the Project data. You can apply to join the GeCIP (https://www.genomicsengland.co.uk/join-a-gecip-domain/). To be eligible for data access, you must have your affiliation verified at an institution that has signed the GeCIP Participation Agreement and have your application to join a valid GeCIP domain accepted by the domain lead or a member of the GeCIP team.

The Incremental Database containing the variant summary statistics at all loci for each data set are available for download at https://doi.org/10.15131/shef.data.9995945 and the gnomAD TSV file generated in this study is available to download at https://doi.org/10.15131/shef.data.9927062. The code used to generate and analyse the IncDBs used in this study is publicly available at https://github.com/tmfreeman400/IncDB_code and as **Supplemental Code**.

## Acknowledgements

Funding support for the analysis and methods development was from the MRC Proximity to Discovery (P2D) Research Scheme and by the NIHR Sheffield Biomedical Research Centre (BRC). The views expressed are those of the author(s) and not necessarily those of the NHS, the NIHR or the Department of Health and Social Care (DHSC). This research was also made possible through access to the data and findings generated by the 100,000 Genomes Project. The 100,000 Genomes Project is managed by Genomics England Limited (a wholly owned company of the Department of Health). The 100,000 Genomes Project is funded by the National Institute for Health Research and NHS England. The Wellcome Trust, Cancer Research UK and the Medical Research Council have also funded research infrastructure. The 100,000 Genomes Project uses data provided by patients and collected by the National Health Service as part of their care and support. The authors gratefully acknowledge Mark Pratt, who initially proposed the concept of the Incremental Database. The authors thank Matt Parker and the Sheffield Bioinformatics Core for their advice on data processing.

## References

Benjamini Y, Speed TP. 2012. Summarizing and correcting the GC content bias in high-throughput sequencing. Nucleic Acids Res 40: e72.

Chen Y-C, Liu T, Yu C-H, Chiang T-Y, Hwang C-C. 2013. Effects of GC bias in next-generation-sequencing data on de novo genome assembly. PLoS One 8: e62856.

Cheung M-S, Down TA, Latorre I, Ahringer J. 2011. Systematic bias in high-throughput sequencing data and its correction by BEADS. Nucleic Acids Res 39: e103.

Cunningham F, Achuthan P, Akanni W, Allen J, Amode MR, Armean IM, Bennett R, Bhai J, Billis K, Boddu S, et al. 2019. Ensembl 2019. Nucleic Acids Res 47: D745–D751.

Goldfeder RL, Priest JR, Zook JM, Grove ME, Waggott D, Wheeler MT, Salit M, Ashley EA. 2016. Medical implications of technical accuracy in genome sequencing. Genome Medicine 8. http://dx.doi.org/10.1186/s13073-016-0269-0.

Hasler J, Strub K. 2007. Survey and Summary: Alu elements as regulators of gene expression. Nucleic Acids Research 35: 1389–1389. http://dx.doi.org/10.1093/nar/gkm044.

Karczewski KJ, Francioli LC, Tiao G, Cummings BB, Alföldi J, Wang Q, Collins RL, Laricchia KM, Ganna A, Birnbaum DP, et al. 2019. Variation across 141,456 human exomes and genomes reveals the spectrum of loss-of-function intolerance across human protein-coding genes. bioRxiv. https://www.biorxiv.org/content/early/2019/01/30/531210.

Karolchik D, Hinrichs AS, Furey TS, Roskin KM, Sugnet CW, Haussler D, Kent WJ. 2004. The UCSC Table Browser data retrieval tool. Nucleic Acids Res 32: D493–6.

Kent WJ, Sugnet CW, Furey TS, Roskin KM, Pringle TH, Zahler AM, Haussler a. D. 2002. The Human Genome Browser at UCSC. Genome Research 12: 996–1006. http://dx.doi.org/10.1101/gr.229102.

King DA, Sifrim A, Fitzgerald TW, Rahbari R, Hobson E, Homfray T, Mansour S, Mehta SG, Shehla M, Tomkins SE, et al. 2017. Detection of structural mosaicism from targeted and whole-genome sequencing data. Genome Res 27: 1704–1714.

Laehnemann D, Borkhardt A, McHardy AC. 2016. Denoising DNA deep sequencing data-high-throughput sequencing errors and their correction. Brief Bioinform 17: 154–179.

Li H. 2013. Aligning sequence reads, clone sequences and assembly contigs with BWA-MEM. http://arxiv.org/abs/1303.3997 (Accessed February 7, 2020).

Li H. 2011. A statistical framework for SNP calling, mutation discovery, association mapping and population genetical parameter estimation from sequencing data. Bioinformatics 27: 2987–2993.

Li H, Handsaker B, Wysoker A, Fennell T, Ruan J, Homer N, Marth G, Abecasis G, Durbin R, 1000 Genome Project Data Processing Subgroup. 2009. The Sequence Alignment/Map format and SAMtools. Bioinformatics 25: 2078–2079.

Mainzer LS, Chapman BA, Hofmann O, Rendon G, Stephens ZD, Jongeneel V. 2015. Validation of Illumina’s Isaac variant calling workflow. bioRxiv. http://dx.doi.org/10.1101/031021.

Martin AR, Williams E, Foulger RE, Leigh S, Daugherty LC, Niblock O, Leong IUS, Smith KR, Gerasimenko O, Haraldsdottir E, et al. 2019. PanelApp crowdsources expert knowledge to establish consensus diagnostic gene panels. Nat Genet. http://dx.doi.org/10.1038/s41588-019-0528-2.

MATLAB, 2019. 9.6 (R2019a). Natick, Massachusetts: The MathWorks Inc.

Ma X, Shao Y, Tian L, Flasch DA, Mulder HL, Edmonson MN, Liu Y, Chen X, Newman S, Nakitandwe J, et al. 2019. Analysis of error profiles in deep next-generation sequencing data. Genome Biol 20: 50.

Raczy C, Petrovski R, Saunders CT, Chorny I, Kruglyak S, Margulies EH, Chuang H-Y, Källberg M, Kumar SA, Liao A, et al. 2013. Isaac: ultra-fast whole-genome secondary analysis on Illumina sequencing platforms. Bioinformatics 29: 2041–2043.

Robinson JT, Thorvaldsdóttir H, Wenger AM, Zehir A, Mesirov JP. 2017. Variant Review with the Integrative Genomics Viewer. Cancer Research 77: e31–e34. http://dx.doi.org/10.1158/0008-5472.can-17-0337.

Ross MG, Russ C, Costello M, Hollinger A, Lennon NJ, Hegarty R, Nusbaum C, Jaffe DB. 2013. Characterizing and measuring bias in sequence data. Genome Biol 14: R51.

Sandmann S, de Graaf AO, Karimi M, van der Reijden BA, Hellström-Lindberg E, Jansen JH, Dugas M. 2017. Evaluating Variant Calling Tools for Non-Matched Next-Generation Sequencing Data. Sci Rep 7: 43169.

Smit, AFA, Hubley, R & Green, P. 2013-2015. RepeatMasker Open-4.0. http://www.repeatmasker.org.

The 1000 Genomes Project Consortium. 2010. A map of human genome variation from population-scale sequencing. Nature 467: 1061–1073.

Vattathil S, Scheet P. 2016. Extensive Hidden Genomic Mosaicism Revealed in Normal Tissue. Am J Hum Genet 98: 571–578.

Wall JD, Tang LF, Zerbe B, Kvale MN, Kwok P-Y, Schaefer C, Risch N. 2014. Estimating genotype error rates from high-coverage next-generation sequence data. Genome Res 24: 1734–1739.

Xiao C, Zook J, Trask S, Sherry S, the Genome-in-a-Bottle Consortium. 2014. Abstract 5328: GIAB: Genome reference material development resources for clinical sequencing. Cancer Research 74: 5328–5328. http://dx.doi.org/10.1158/1538-7445.am2014-5328.

Yizhak K, Aguet F, Kim J, Hess JM, Kübler K, Grimsby J, Frazer R, Zhang H, Haradhvala NJ, Rosebrock D, et al. 2019. RNA sequence analysis reveals macroscopic somatic clonal expansion across normal tissues. Science 364. http://dx.doi.org/10.1126/science.aaw0726.

Zook JM, Chapman B, Wang J, Mittelman D, Hofmann O, Hide W, Salit M. 2014. Integrating human sequence data sets provides a resource of benchmark SNP and indel genotype calls. Nat Biotechnol 32: 246–251.

Zook JM, McDaniel J, Olson ND, Wagner J, Parikh H, Heaton H, Irvine SA, Trigg L, Truty R, McLean CY, et al. 2019. An open resource for accurately benchmarking small variant and reference calls. Nat Biotechnol 37: 561–566.

