## Supplemental Materials for "Genomic loci susceptible to systematic sequencing bias in clinical whole genomes"

**Contents:** This document lists all of the supplemental materials provided with this publication for download, along with captions describing each file. These include:

Supplemental Code

Supplemental Data S1-2

Supplemental Tables S1-2

Supplemental Figures S1-12

**Supplemental Code**: All of the code we used to generate Incremental Databases is available in the zip file available for download in the supplemental materials, along with instructions for how to use this. This code is also publicly available in the GitHub repository at <https://github.com/tmfreeman400/IncDB_code> .

**Supplemental Data S1** - List of autosomal genes associated with hereditary spastic paraplegia from the PanelApp gene panel, ranked in descending order by suspect loci percentage. Files A, B, C refer to data sets 1, 2, 3 respectively.

**Supplemental Data S2** - List of autosomal genes associated with genetic epilepsy syndromes from the PanelApp gene panel, ranked in descending order by suspect loci percentage. Files A, B, C refer to data sets 1, 2, 3 respectively.

|  | **Number of suspect loci**  **(×1,000,000)** | **Number of non-suspect loci**  **(×1,000,000)** | **Odds ratio** | **Fisher Exact Test p-value** |
| --- | --- | --- | --- | --- |
| **Inside GC-rich regions** | **1.59** | **14.7** | **8.19** | **1.10×10‾³²¹** |
| **Outside GC-rich regions** | **34.8** | **2630** |  |  |

**Supplemental Table S1 Extract (Full Supplemental Table S1 available for download with this paper)**: A Fisher Exact test contingency table for suspect locus enrichment within GC-rich regions, across all autosomal chromosomes (from data set 1). The full contingency table for all regions across data sets 1-3 is available in Supplemental_Table_S1.xlsx, showing these statistics for all 18 types of region examined in this study, with the exception of the p-value, since this was extremely low in all cases (the largest p-value was 8.69×10‾⁴¹ for the enrichment of suspect loci in the intellectual disability gene panel in data set 2, which was still highly statistically significant).

**Supplemental Table S2:** A table showing the distribution of patient ethnicities in data sets 2-5 (this data was not available for data set 1). The number of patients for each of the 16 ethnic groups is listed along with the total number of patients for the data set. All categories are distinct and no patient is in more than one category.

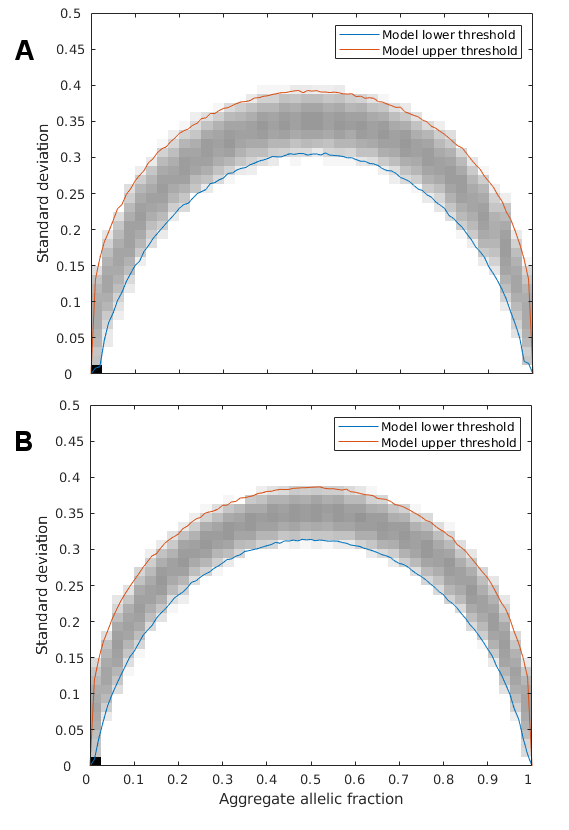

**Supplemental Figure S1:** Monte Carlo random sampling was used to generate the Hardy-Weinberg expected range of standard deviation at each allelic fraction with 1000 repetitions at each. The standard deviation and aggregate allelic fraction for these simulated values are plotted on the above density plot. The maximum and minimum boundaries of these values are marked in red and blue respectively and represent the upper and lower 99.9% confidence interval for all autosomal chromosomes with a population of 150 patients (A), corresponding to data set 1, or 215 patients (B), corresponding to data sets 2 and 3.

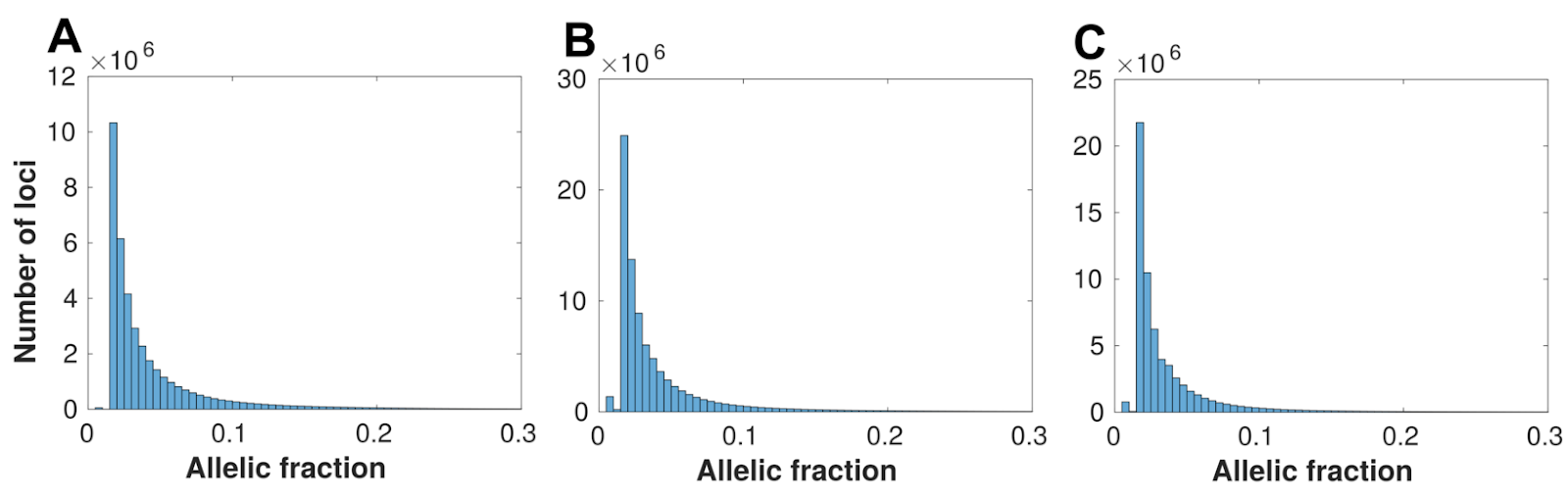

**Supplemental Figure S2**: Histograms showing the distribution of allelic fractions observed at suspect loci in data sets 1-3 respectively (A-C), using the default suspect locus classification parameters for this study (confidence interval=99.9%, assumed error rate=0.01). Each bin has a width of 0.005 allelic fraction and the scale is cut off at allelic fraction=0.3 because the number of suspect loci decreases to near-zero. The allele fractions of the systematic errors were generally very small and heavily right-skewed (median= 2.82%, 2.61%, 2.41%; 95% confidence interval = 1.54-18.84%, 1.51-20.59%, 1.51-17.70% for data sets 1-3 respectively).

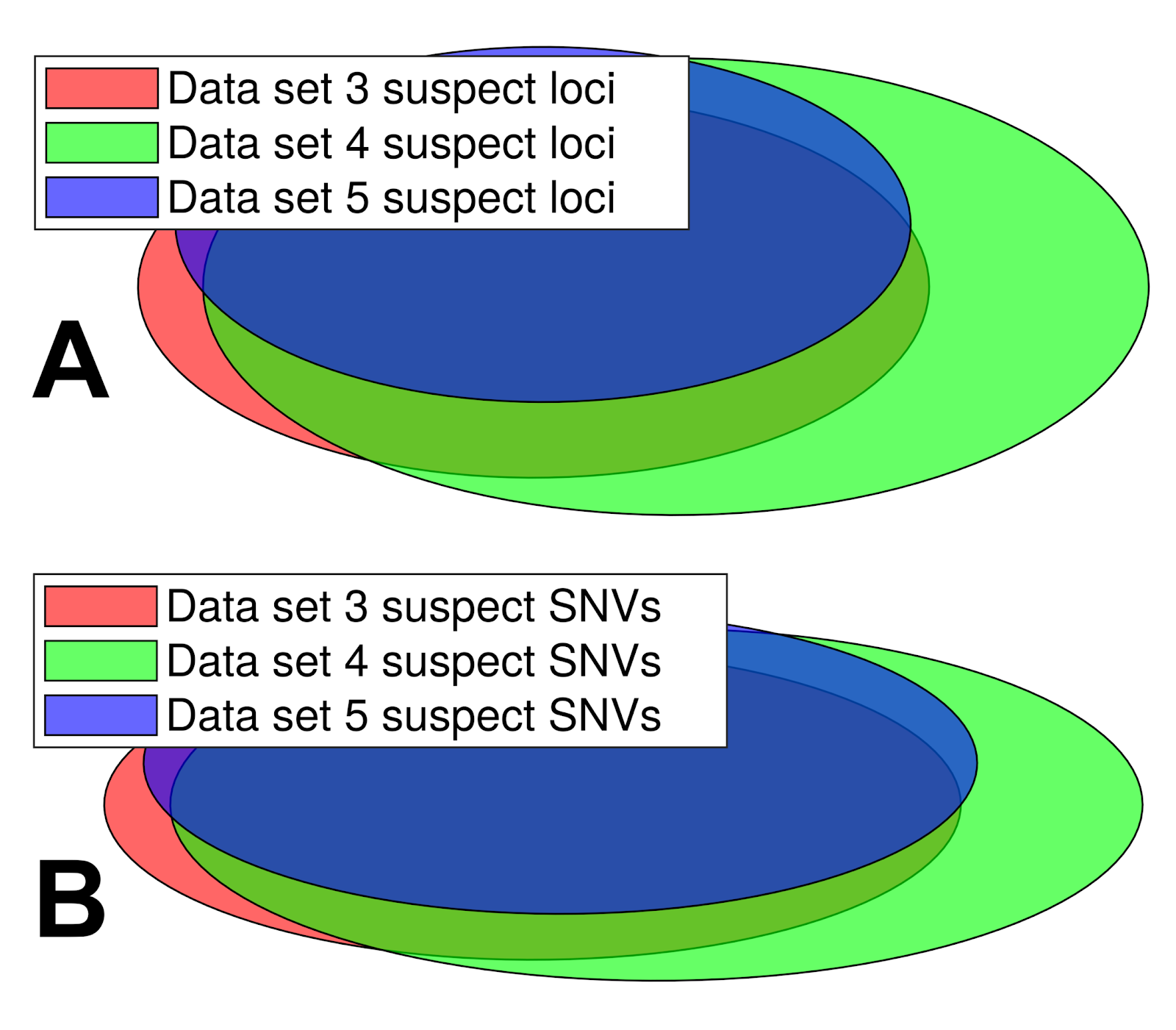

**Supplemental Figure S3**: Venn diagrams showing the overlap in data sets 3-5 of (**A**) all unique suspect loci and (**B**) called SNVs in NA12878 annotated as suspect, with shaded areas proportional to the numbers and overlap of suspect loci and suspect SNVs in each data set. All 3 of these data sets use exactly the same technical conditions, but share no patients in common. All the patients used have the same distribution of clinical attributes but cohort 5 are on average an older group to explore the possibility of age-related mosaicism impacting suspect loci detected. These 3 IncDBs show considerable suspect loci overlap (mean overlap=65.7%) and suspect SNV overlap (mean overlap=71.9%) between each other, although there are some differences likely due to private mutations. The higher degree of overlap between data sets 3 and 5 than between data sets 3 and 4, as well as the lower numbers of suspect loci and suspect SNVs in data set 5 than data set 4 suggest that age-related mosaicism is unlikely to impact on suspect locus annotation.

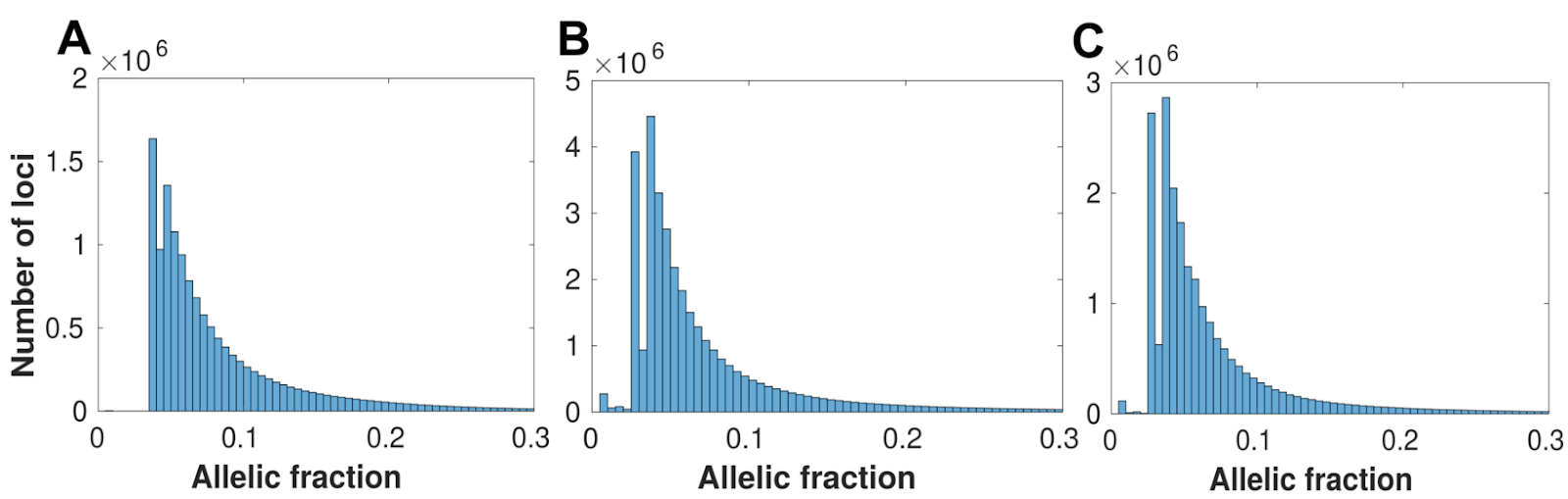

**Supplemental Figure S4**: Histograms showing the distribution of allelic fractions observed at suspect loci in data sets 1-3 respectively (A-C), using an alternative, stricter confidence threshold for suspect locus classification than the default parameters used in this study (confidence interval=99.999%, assumed error rate=0.01). Each bin has a width of 0.005 allelic fraction and the scale is cut off at allelic fraction=0.3 because the number of suspect loci decreases to near-zero. Compared to Supplemental Fig. S2, the number of suspect loci at lower allelic fractions (<3%) drops off sharply since the systematic error rate is unlikely to be significant at such a low effect size, at this confidence level.

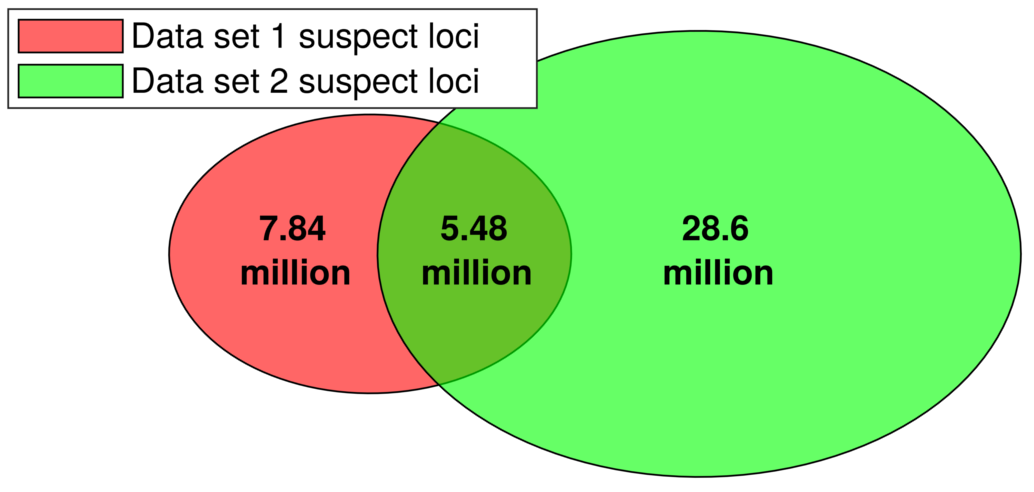

**Supplemental Figure S5**: Venn diagram showing the overlap of all unique suspect loci between data sets 1 and 2, using an alternative, stricter confidence threshold for suspect locus classification (confidence interval=99.999%, assumed error rate=0.01). Data set 3 used the GRCh38 reference, so it was not included in the Venn diagram. Applying this alternative threshold filtered out a very high proportion (99.98/91.06/92.68% in data sets 1, 2 and 3 respectively) of the suspect loci that had allelic fractions <0.03, since fewer low allelic fraction loci exhibited a standard deviation lower than the 99.999% confidence threshold. In contrast, a much lower proportion (25.95/17.74/23.40% in data sets 1, 2 and 3 respectively) of the suspect loci with larger systematic biases (allelic fractions>0.03) were filtered out at the 99.999% confidence threshold. The percentage overlap of suspect loci between data sets 1 and 2 decreases slightly when the more conservative threshold (99.999% confidence interval) is used for defining suspect loci (from 47.9/21.7% with the default threshold (Fig. 2D) to 41.1/16.1% in this figure, in data sets 1/2 respectively).

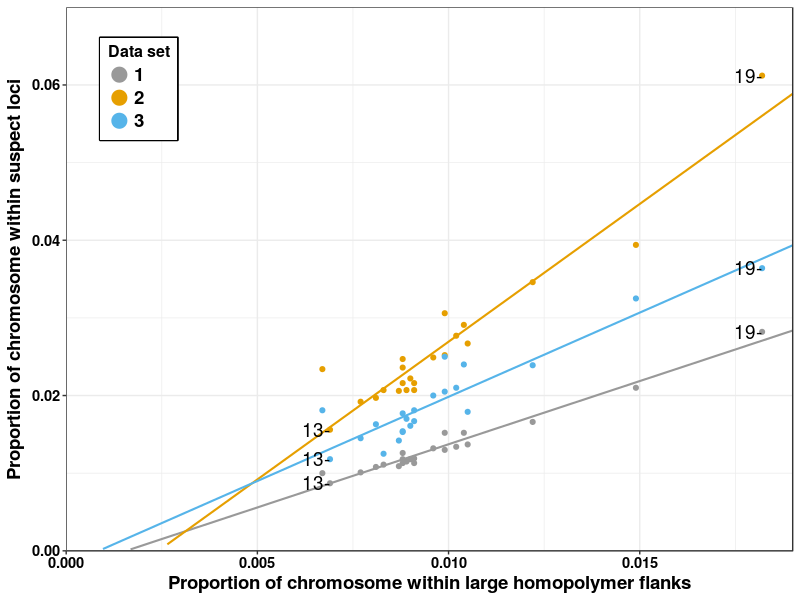

**Supplemental Figure S6**: Proportion of chromosomal positions with at least one suspect locus vs. proportion of chromosome within large homopolymer flanks, with linear line of best fit. There is a strong linear correlation between the proportion of unique suspect loci in each chromosome and the proportion of loci in large homopolymer flanks, indicating that these regions are a key contributor to systematic sequencing and/or alignment errors. Chromosomes 13 and 19 are labelled as the chromosomes with the lowest and highest levels of suspect loci respectively in all 3 data sets.

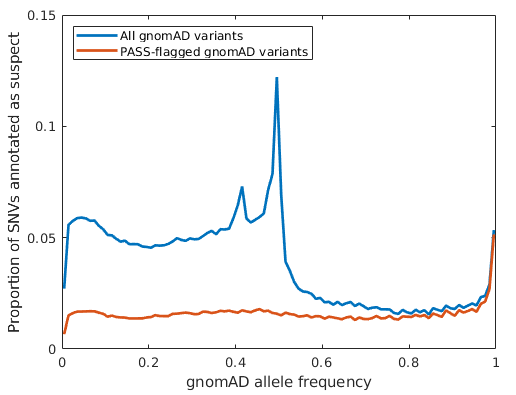

**Supplemental Figure S7**: Proportion of autosomal gnomAD SNVs annotated as suspect at different gnomAD allele frequencies in data set 1. A large proportion of the low allelic frequency gnomAD variants annotated as suspect did not pass gnomAD’s quality control, but ~1.5% of SNVs that passed gnomAD’s quality control checks were annotated as suspect across most allele frequencies, suggesting that systematic biases are prevalent in gnomAD’s called SNVs, albeit at a low rate.

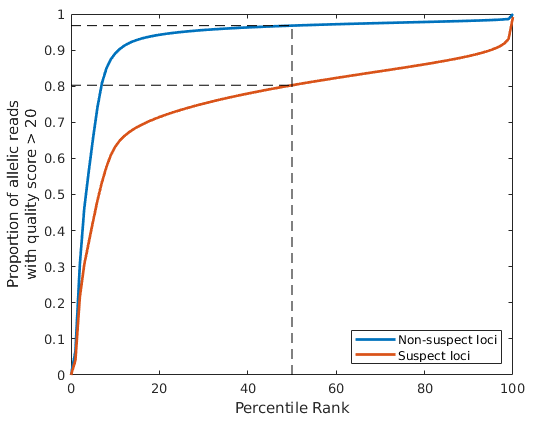

**Supplemental Figure S8**: The proportion of allelic reads that had quality scores > 20 (for both sequencing and mapping), plotted against the percentile ranks for both non-suspect loci (blue) and suspect loci (red) respectively using this measure, including reads for all locus/allele combinations for Chromosome 1 in data set 1 (Personalis Inc.). Non-suspect loci had significantly higher proportions of high quality reads, with >90% of reads having quality scores >20 in ~90% of non-suspect loci/allele combinations vs. ~5% of suspect loci/allele combinations.

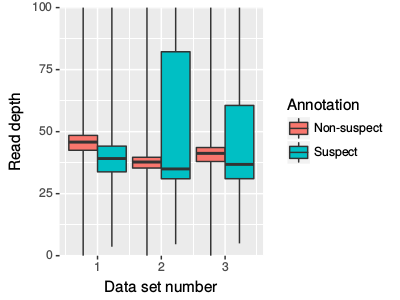

**Supplemental Figure S9**: Boxplots showing the distribution (quartiles) of read depths observed across suspect (cyan) and non-suspect (red) loci positions in data sets 1-3 respectively. The maximum read depth values for the whiskers were 5,304/6,165/6,318 for non-suspect loci and 5,195/9,093/7,764 for suspect loci in data sets 1-3 respectively, but the figure y limit was kept at a read depth of 100 for clarity over the main range of read depths. Two-Sample Kolmogorov-Smirnov tests confirmed that there was a different distribution of read depth between genomic positions with and without suspect loci (D=0.4289, 0.2885, 0.3079 and p<10‾¹⁵ for data sets 1, 2, 3 respectively), but there was no clear correlation between read depth and likelihood of a position having suspect loci. While the lower quartile and median values for read depth were slightly lower for unique suspect loci, for data sets 2 and 3 the upper quartiles were significantly higher at unique suspect loci.

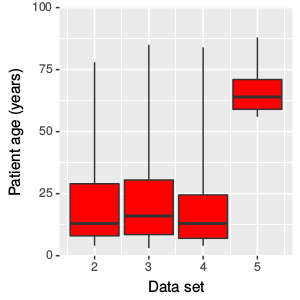

**Supplemental Figure S10**: Boxplots showing the distribution (quartiles) of patient ages in data sets 2-5. This information was not available for data set 1, but we were told that the patients in that data set were all trios including two parents and an infant, so the age spread for data set 1 was ⅓ very young patients and ⅔ medium age patients. For data set 5 we intentionally selected older patients from the 100,000 Genomes Project than data sets 2-4, to identify if age had an effect on suspect locus annotation, but did not find any evidence for this.

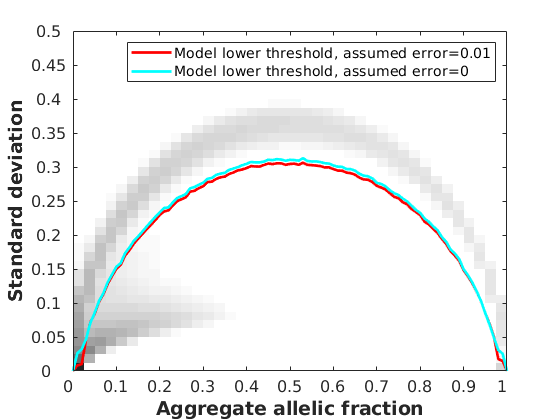

**Supplemental Figure S11**: Density plot showing the distribution of aggregate allelic fraction and standard deviation values for all autosomal chromosomes in the Incremental Database for data set 1, alongside the maximum standard deviation limits used to annotate suspect loci assuming error rates of 0.01 (red) and 0 (cyan) respectively, which were obtained using Monte Carlo simulations at a 99.9% confidence interval. The greatest difference between these thresholds can be observed at aggregate allelic fractions of <0.03, which results in an increase in suspect loci from 38.7 million to 137.9 million in data set 1. However, loci at these low allelic fractions are rarely called due to low number of supporting reads, and there is only a small resulting increase (200,265 to 214,522) in the number of suspect loci with called SNVs when we reduce the error rate.

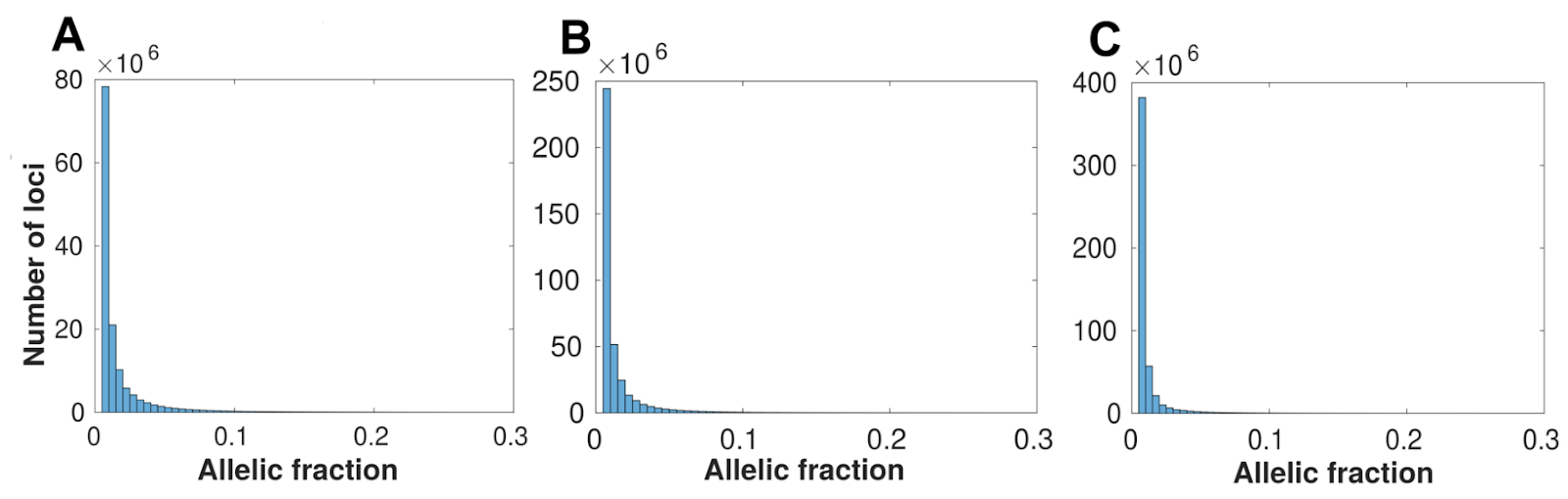

**Supplemental Figure S12**: Histograms showing the distribution of allelic fractions observed at suspect loci in data sets 1-3 respectively (A-C), using an alternative, less conservative threshold for suspect locus classification than the default parameters used in this study (confidence interval=99.9%, assumed error rate=0). Each bin has a width of 0.005 allelic fraction and the scale is cut off at allelic fraction=0.3 because the number of suspect loci decreases to near-zero. Compared to Supplemental Fig. S2, the number of suspect loci at very low allelic fractions (0.5-1.5%) is much higher since decreasing the assumed error rate to zero raises the maximum standard deviation limit at very low allelic fractions greatly, causing many more of these loci to fall under it, but does not greatly affect the numbers of suspect loci annotated at >3% allelic fraction.
