## Supplemental Code for "Genomic loci susceptible to systematic sequencing bias in clinical whole genomes": build_empty_matrices.R

```
#!/usr/bin/env Rscript
args = commandArgs(trailingOnly = TRUE)

#This script creates empty files of zeros for chromosomes 1-22
#The chromosome number should be passed to this script as the first argument (e.g. 1, 2, 3, etc.)
#The genome build should be passed to this script as the second argument (37 or 38)
#The project directory should be passed to this script as the third argument

#Set working directory to the same directory used in the 2nd line of structure_directories.sh
setwd(args[3])

for(i in as.numeric(args[1])){
chrnum = i

#Get lenngths of all chromosomes for their genome build
chrlengths37 = c(249250621, 243199373, 198022430, 191154276,
               180915260, 171115067, 159138663, 146364022,
               141213431, 135534747, 135006516, 133851895,
               115169878, 107349540, 102531392, 90354753,
               81195210, 78077248, 59128983, 63025520,
               48129895, 51304566)

chrlengths38 = c(248956422, 242193529, 198295559, 190214555,
                 181538259, 170805979, 159345973, 145138636,
                 138394717, 133797422, 135086622, 133275309,
                 114364328, 107043718, 101991189, 90338345,
                 83257441, 80373285, 58617616, 64444167,
                 46709983, 50818468)

chrlength37 = chrlengths37[chrnum]
chrlength38 = chrlengths38[chrnum]

if(as.numeric(args[2]==37)){
zeromatrix37 = matrix(0,chrlength37,4)
filename37 = paste0('Empty_IncDB37_', chrnum)
write.table(zeromatrix37, file = filename37, row.names = FALSE, col.names = FALSE, sep = '\t')
}

if(as.numeric(args[2]==38)){
zeromatrix38 = matrix(0,chrlength38,4)
filename38 = paste0('Empty_IncDB38_', chrnum)
write.table(zeromatrix38, file = filename38, row.names = FALSE, col.names = FALSE, sep = '\t')
}

}
```
