## Supplemental Code for "Genomic loci susceptible to systematic sequencing bias in clinical whole genomes": get_af.R

```
#!/usr/bin/env Rscript
args = commandArgs(trailingOnly = TRUE)

#get_af function takes patient number as the first argument and
#chromosome number as the second argument

#Edit the filepaths in this section to the corresponding filepaths on your system so that the code functions correctly
pat_filepaths  <- 'path_to_textfile_listing_patient_bam_filepaths' #REPLACE this with the filepath to your list
project_directory <- 'path_to_project_directories' #REPLACE this with the filepath to the directories created by the build_directories.R command
build_number <- 38 #Replace with 37 if you are using build 37

#NO EDITS SHOULD BE REQUIRED AFTER THIS POINT
args[3] <- build_number

project_directory <- paste0(project_directory, '/')
directories <- unlist(strsplit(scan(pat_filepaths, what = 'character')[as.numeric(args[2])], split='/'))
finalsplit <- paste0('_', directories[length(directories)-1], '_') #Name of the last sub-directory in which bam files are located

#NO EDITS SHOULD BE REQUIRED AFTER THIS POINT

#Create solo DB row-by-row by adding base counts where indicated
#And saving separate record for each patient
patnum_in_list = as.numeric(args[1])
chrnum = as.numeric(args[2])

#Set working directory to corresponding chromosome
setwd(paste0(project_directory, 'chr', chrnum))

#Get patient names with available base counts from working directory
gpat=paste0('chr',chrnum,'chunk')
patnames=grep(pattern = gpat, x = dir(), value = TRUE)
stems=unlist(strsplit(patnames, split = gpat))[seq(from=1, by=2, to=(2*length(patnames)-1))]
uniq_stems=unique(stems)
pat_fragments = unlist(strsplit(unlist(scan(pat_filepaths, what = 'character'))[patnum_in_list], split = '/'))
patnum = grep(pattern = pat_fragments[length(pat_fragments)], x = uniq_stems)

#Check if soloDB already exists and stop job if so.
#Comment below 11 lines to remove this step
patientsstem <- paste0(unlist(strsplit(paste0(unlist(strsplit(uniq_stems[patnum], split = finalsplit)))[2], split = '.bam')))[1]
if(length(grep('solo', system('ls', intern = TRUE)))!=0){
  filesizes <- system(paste0("ls -l ", project_directory, "/chr", chrnum, "/*soloDB | awk '{print $5}' "), intern = TRUE)
  allfiles  <- system(paste0("ls -l ", project_directory, "/chr", chrnum, "/*soloDB | awk '{print $9}' "), intern = TRUE)
  zerofiles <- allfiles[which(filesizes=='0')]

  #Check if file already exists with non-zero size
  if(length(grep(patientsstem, allfiles))>0){
    #Do not repeat if so
    if(length(grep(patientsstem, zerofiles))==0){stop()}
  }
}

#
num_pats=length(uniq_stems)
last_chunk_num <- c(5,5,4,4,4,4,4,3,3,3,3,3,3,3,3,2,2,2,2,2,1,2)[chrnum]

for(patno in patnum){
  
  #Select chunk
  
  for(chunknum in 1:last_chunk_num){
  
  #Generate DBtofill for this chunk- either 50M rows or smaller if last chunk
  
  for(chrnum in as.numeric(args[2])){
    
    #Get lenngths of all chromosomes for their genome build
    chrlengths37 = c(249250621, 243199373, 198022430, 191154276,
                     180915260, 171115067, 159138663, 146364022,
                     141213431, 135534747, 135006516, 133851895,
                     115169878, 107349540, 102531392, 90354753,
                     81195210, 78077248, 59128983, 63025520,
                     48129895, 51304566)
    
    chrlengths38 = c(248956422, 242193529, 198295559, 190214555,
                     181538259, 170805979, 159345973, 145138636,
                     138394717, 133797422, 135086622, 133275309,
                     114364328, 107043718, 101991189, 90338345,
                     83257441, 80373285, 58617616, 64444167,
                     46709983, 50818468)
    
    chrlength37 = chrlengths37[chrnum]
    chrlength38 = chrlengths38[chrnum]
    
    if(as.numeric(args[3]==37)){maxchrlength = chrlength37}
    if(as.numeric(args[3]==38)){maxchrlength = chrlength38}
    
    #Get chunklength from maxchrlength and chunknum
    if(chunknum<last_chunk_num){chunklength <- 50000000}
    if(chunknum==last_chunk_num){chunklength <- maxchrlength-(chunknum-1)*50000000}
    
    DBtofill = matrix(0,chunklength,4)
    
  }
  
  #Fill DBtofill looping through each allele
  
  for(basetype in 1:4){
    baseallele = c('A','C','G','T')[basetype]
    
    #Read in data from chunk
    filler=read.table(paste0(uniq_stems[patno], gpat, chunknum, '_', baseallele,'counts.txt'),
                      header = FALSE, as.is = TRUE)
    
    #Write count values to matrix at corresponding rows
    rowindices = as.numeric(filler[,2])
    rowcounts = as.numeric(filler[,1])
    number_of_filling_rows = length(rowindices)
    DBtofill[rowindices, basetype] <- rowcounts

  }
  
  #Divide rows of matrix by total to convert absolute counts to allelic fractions
  DBtofill <- DBtofill/rowSums(DBtofill)
  
  is.nan.data.frame <- function(x)
    do.call(cbind, lapply(x, is.nan))
  
  #Replace NA with 0 since this corresponds to zero AF for all alleles
  DBtofill[is.nan(DBtofill)] <- 0
  
  #Save results for single patient
  write.table(DBtofill, file = paste0(uniq_stems[patno], 'chr', chrnum, '_soloDBchunk', chunknum), row.names = FALSE, col.names = FALSE, sep = '\t')
  
  }

  #cat chunks together into finished soloDB and rm chunks
  names2cat <- paste0(uniq_stems[patno], 'chr', chrnum, '_soloDBchunk*')
  cattedoutput <- paste0(uniq_stems[patno], 'chr', chrnum, '_soloDB')
  systemcommand = paste0("cat ", names2cat, " > ", cattedoutput)
  system(systemcommand, wait = TRUE)
  
  system(paste0("rm ", names2cat), wait = TRUE)
  }

#For next step run get_mean_af.sh and get_sd_af.sh
```
