## Supplemental Code for "Genomic loci susceptible to systematic sequencing bias in clinical whole genomes": get_mean_af.sh

```
#!/bin/bash

#First input is filepath to project directory
#Second input is chromosome number
var1=$1
var2=$2
c="/chr"

cd $var1$c$var2

#Get number of 1million bp chunks
solosize=`wc *soloDB | awk '{print $1}' | head -n 1`
numchunks=`echo $((1+$(($solosize/1000000))))`
echo $numchunks

currentchunk=1
while [[ $currentchunk -le $numchunks ]]; do
maxco=`echo $(($currentchunk*1000000))`
#Generate chunks
tag="_mchunk"
for file in *soloDB; do
awk -v maxc=$maxco 'NR==maxc-999999, NR==maxc-1; NR==maxc {print; exit}' $file > $file$tag$currentchunk
done;
#Do for loop for mawk code over chunks to calculate mean (aggregate) allelic frequencies
mawk '{a[FNR]+=$1; b[FNR]++;
d[FNR]+=$2; e[FNR]++;
g[FNR]+=$3; h[FNR]++;
j[FNR]+=$4; k[FNR]++;}
	END{
	  for(i=1;i<=FNR;i++)
	    print a[i]/b[i],
                  d[i]/e[i],
                  g[i]/h[i],
                  j[i]/k[i]
 ;
   }' *soloDB$tag$currentchunk > meanAF$tag$currentchunk.txt
currentchunk=`echo $((1+$currentchunk))`
done

#Merge chunks into output
currentchunk=1
cat meanAF_mchunk1.txt > meanAF.txt
while [[ $currentchunk -lt $numchunks ]]; do
nextchunk=`echo $((1+$currentchunk))`
cat meanAF.txt meanAF_mchunk$nextchunk.txt > tempmeanAF.txt
cat tempmeanAF.txt > meanAF.txt
currentchunk=`echo $nextchunk`
done

#Delete chunks and temporary files
rm tempmeanAF.txt
rm *meanAF_mchunk?.txt
rm *meanAF_mchunk??.txt
rm *meanAF_mchunk???.txt
rm *mchunk?
rm *mchunk??
rm *mchunk???
```
