## Supplemental Code for "Genomic loci susceptible to systematic sequencing bias in clinical whole genomes": get_sd_af.sh

```
#!/bin/bash

#First input is filepath to project directory
#Second input is chromosome number
var1=$1
var2=$2
c="/chr"

cd $var1$c$var2

#Get number of 1million bp chunks
solosize=`wc *soloDB | awk '{print $1}' | head -n 1`
numchunks=`echo $((1+$(($solosize/1000000))))`
echo $numchunks

currentchunk=1
while [[ $currentchunk -le $numchunks ]]; do
maxco=`echo $(($currentchunk*1000000))`
#Generate chunks
tag="_schunk"
for file in *soloDB; do
awk -v maxc=$maxco 'NR==maxc-999999, NR==maxc-1; NR==maxc {print; exit}' $file > $file$tag$currentchunk
done;
#Do for loop for mawk code over chunks to calculate standard deviation between allelic frequencies
mawk '{a[FNR]+=$1; b[FNR]++; c[FNR]+=($1)^2;
d[FNR]+=$2; e[FNR]++; f[FNR]+=($2)^2;
g[FNR]+=$3; h[FNR]++; i2[FNR]+=($3)^2;
j[FNR]+=$4; k[FNR]++; l[FNR]+=($4)^2;}
	END{
	  for(i=1;i<=FNR;i++)
	    print sqrt((c[i]-a[i]*a[i]/b[i])/(b[i]-1)),
                  sqrt((f[i]-d[i]*d[i]/e[i])/(e[i]-1)),
                  sqrt((i2[i]-g[i]*g[i]/h[i])/(h[i]-1)),
                  sqrt((l[i]-j[i]*j[i]/k[i])/(k[i]-1))
 ;
   }' *soloDB$tag$currentchunk > sdAF$tag$currentchunk.txt
currentchunk=`echo $((1+$currentchunk))`
done

#Merge chunks into output
currentchunk=1
cat sdAF_schunk1.txt > sdAF.txt
while [[ $currentchunk -lt $numchunks ]]; do
nextchunk=`echo $((1+$currentchunk))`
cat sdAF.txt sdAF_schunk$nextchunk.txt > tempsdAF.txt
cat tempsdAF.txt > sdAF.txt
currentchunk=`echo $nextchunk`
done

#Delete chunks and temporary files
rm tempsdAF.txt
rm *sdAF_schunk?.txt
rm *sdAF_schunk??.txt
rm *sdAF_schunk???.txt
rm *schunk?
rm *schunk??
rm *schunk???
```
