## Supplemental Code for "Genomic loci susceptible to systematic sequencing bias in clinical whole genomes": mpileup_read_count.R

```
#!/usr/bin/env Rscript
args = commandArgs(trailingOnly = TRUE)

#ATTENTION: Need to module load samtools prior to running on headnode
#You might need to edit this command for your system so that samtools is available
system('module load samtools/1.9')

#Edit the filepaths in this section to the corresponding filepaths on your system so that the code functions correctly
pat_filepaths  <- 'path_to_textfile_listing_patient_bam_filepaths' #REPLACE this with the filepath to your list
project_directory <- paste0('path_to_project_directories','/') #REPLACE this with the filepath to the directories created by the build_directories.R command
build_number <- 38 #Replace with 37 if you are using build 37

#NO EDITS SHOULD BE REQUIRED AFTER THIS POINT

#R script to get all allelic counts ACGT for specified patient (first argument) for specified chr (second argument)

#Extract script inputs for patient and chromosome number for use in script
#First argument is the row number of a text file that corresponds to the row 
#that contains the filepath of the bam file referring to that patient
patid <- patientid <- as.numeric(args[1]);
#Second argument is the number of the chromosome
chrinput <- as.numeric(args[2]);

#Define function to run for patient's bam file
mpileup_function <- function(patient, patid, chrinput){
  
  #Store patient ID values
  patientid <- patid;
  
  #Run series of commands for specific chromosome
  for(chrnum in chrinput){
    
    #Use representative non-cancer patient sample (recommended to be 100 patients or more if possible)
    chosen <- scan(pat_filepaths, what = 'character')
    patient_filepath <- chosen[patientid] #This is the filepath to the bam file you will be counting reads in
    
    #Get string corresponding to chromosome name in the bam file, based on the number
    chrlist <- c('1', '2', '3', '4', '5', '6', '7', '8', '9', '10',
                 '11','12','13','14','15','16','17','18','19', '20',
                 '21','22', 'X', 'Y', 'MT')
    chrstr <- chrlist[chrnum]
    
    #Check that file doesn't already exist, and if so,
    #whether this script previously terminated early giving a file size of zero
    filesizes <- system(paste0("ls -l ", project_directory, "/chr", chrstr, "/* | awk '{print $5}' "), intern = TRUE)
    allfiles  <- system(paste0("ls -l ", project_directory, "/chr", chrstr, "/* | awk '{print $9}' "), intern = TRUE)
    zerofiles <- allfiles[which(filesizes=='0')]
    
    #Get file name stem that would correspond to this file for following checks
    num_slashes <- length(paste0(unlist(strsplit(chosen[patientid], split = '/'))))-1
    patientsstem <- paste0(unlist(strsplit(paste0(unlist(strsplit(chosen[patientid], split = '/')))[(1+num_slashes)], split = '.bam')))[1]
    
    #Check if file already exists. If so:
    if(length(grep(patientsstem, allfiles))>0){
      #If file already exists and has non-zero size, do not repeat
      if(length(grep(patientsstem, zerofiles))==0){next()}
      #If not, above line is skipped
    }
    
    chrlist <- c('1', '2', '3', '4', '5', '6', '7', '8', '9', '10',
                 '11','12','13','14','15','16','17','18','19', '20',
                 '21','22', 'X', 'Y', 'MT')
    
    p <- patient
    
    unlisted <- unlist(strsplit(p, split = '/'))
    
    p2stem <-  paste0(unlisted[2:length(unlisted)],collapse = '/')
    p3stem <-  paste0(unlisted[2:length(unlisted)],collapse = '_')
    p2 <- chosen[patientid]
    
    #Define number of chunks based off of chromosome size (same for build 37 and 38)
    last_chunk_num <- c(5,5,4,4,4,4,4,3,3,3,3,3,3,3,3,2,2,2,2,2,1,2)[chrnum]
    
    #Write files in 50 million bp chunks of chromosome at a time 
    for(chunk_number in 1:last_chunk_num){
      
      #Check if chunk file already exists. If so:
      if(length(intersect(grep(patientsstem, allfiles), grep(paste0('chunk', chunk_number), allfiles)))!=0){
        #If chunk file already exists and has non-zero size, go to next value in loop,
        #since chunk file has already been created and we do not want to overwrite it.
        if(length(intersect(grep(patientsstem, zerofiles), grep(paste0('chunk', chunk_number), zerofiles)))==0){next()}
        #If not, above line is skipped and chunk file is generated as detailed below.
      }
      
      startcoord <- ((chunk_number-1)*50000000+1)
      endcoord <- chunk_number*50000000
      
      #Run mpileup commands for specified chromosome
      for(chromnum in chrlist[chrnum]){
        
        #Loop over alleles A,C,G,T
        for(allele_letter in c('A', 'C', 'G', 'T')){
          
          #print(p2)
          #print(chromnum)
          #print(allele_letter)
          
          #Write file path to use for chunk files generated
          chunk_out_path <- paste0(project_directory, "chr", chrstr, "/", p3stem ,"_chr", chromnum, "chunk", chunk_number,"_", allele_letter ,"counts.txt")
          
          # chr might not be needed after -r in "samtools mpileup -r chr", depending upon whether
          #chromosomes are labelled as 1,2,3 etc. (omit) or chr1, chr2, chr3, etc. (include chr)
          #Change temporary directory for command if running out of space
          systemcommand <- paste0("export TMPDIR=", project_directory,"; samtools mpileup -r chr", chromnum ,
                                  ":",as.integer(startcoord),"-",as.integer(endcoord)," -Q 0 -x -a  ", p2 ,
                                  " | awk '{print $5}' | grep -i '", allele_letter ,
                                  "' -o -n | cut -d : -f 1 | sort -n | uniq -c > ", chunk_out_path)
          
          system(systemcommand, wait = TRUE)
          
        }}
      
      #Try again with different command (same except for 'chr' omission after -r in "samtools mpileup -r chr") if files are still empty
      filesizes <- system(paste0("ls -l ", project_directory, "/chr", chrstr, "/* | awk '{print $5}' "), intern = TRUE)
      allfiles  <- system(paste0("ls -l ", project_directory, "/chr", chrstr, "/* | awk '{print $9}' "), intern = TRUE)
      zerofiles <- allfiles[which(filesizes=='0')]
      
      #Check if chunk file already exists. If so:
      if(length(intersect(grep(patientsstem, allfiles), grep(paste0('chunk', chunk_number), allfiles)))!=0){
        #If chunk file already exists and has non-zero size, go to next value in loop,
        #since chunk file has already been created and we do not want to overwrite it.
        if(length(intersect(grep(patientsstem, zerofiles), grep(paste0('chunk', chunk_number), zerofiles)))==0){next()}
        #If not, above line is skipped and chunk file is generated as detailed below.
      }
      
      for(chromnum in chrlist[chrnum]){
        
        #Loop over alleles A,C,G,T
        for(allele_letter in c('A', 'C', 'G', 'T')){
          
          #print(p2)
          #print(chromnum)
          #print(allele_letter)
          
          #Write file path to use for chunk files generated
          chunk_out_path <- paste0(project_directory, "chr", chrstr, "/", p3stem ,"_chr", chromnum, "chunk", chunk_number,"_", allele_letter ,"counts.txt")
          
          systemcommand <- paste0("export TMPDIR=", project_directory,"; samtools mpileup -r", chromnum ,
                                  ":",as.integer(startcoord),"-",as.integer(endcoord)," -Q 0 -x -a  ", p2 ,
                                  " | awk '{print $5}' | grep -i '", allele_letter ,
                                  "' -o -n | cut -d : -f 1 | sort -n | uniq -c > ", chunk_out_path)
          
          system(systemcommand, wait = TRUE)
          
        }}
      
      #If chunk file still has zero size, stop execution, there is an error
      filesizes <- system(paste0("ls -l ", project_directory, "/chr", chrstr, "/* | awk '{print $5}' "), intern = TRUE)
      allfiles  <- system(paste0("ls -l ", project_directory, "/chr", chrstr, "/* | awk '{print $9}' "), intern = TRUE)
      zerofiles <- allfiles[which(filesizes=='0')]
      #Check if chunk file already exists. If so:
      if(length(intersect(grep(patientsstem, allfiles), grep(paste0('chunk', chunk_number), allfiles)))!=0){
        #If chunk file already exists and has zero size, stop execution
        if(length(intersect(grep(patientsstem, zerofiles), grep(paste0('chunk', chunk_number), zerofiles)))!=0){stop()}
        #If not, above line is skipped
      }
      
    }
  }}#End of mpileup function

#Set arguments for mpileup function
chosen <- scan(pat_filepaths, what = 'character')
patient_filepath <- chosen[patientid] #This is the filepath to the bam file you will be counting reads in

mpileup_function(patient_filepath, patientid, chrinput)

#For the next step (generating soloDBs) run get_af.R
```
