## Supplemental Code for "Genomic loci susceptible to systematic sequencing bias in clinical whole genomes": structure_directories.sh

```
#Select directory in which to run project (should have very large filespace available ~1TB)
#Input is filepath to project directory
var1=$1

cd $var1

#Create directories and subdirectories for all chromosomes
for i in {1..22}
do
  mkdir chr$i
done

#Run build_empty_matrices after this step
```
