## Supplemental Code for "Genomic loci susceptible to systematic sequencing bias in clinical whole genomes": Workflow_for_IncDB_code

**README file for running code to generate IncDB from .bam files**

Dependencies:

-Samtools (we used version 1.9) required for the mpileup command
-R version 3.4.4 (2018-03-15) or later

Steps:

1)
-Ensure dependencies are loaded.

2)
-Run structure_directories.sh in the directory you want to use for carrying out these steps. We recommend creating an empty directory for this.
Pass the desired project directory filepath to this script as the only argument.
This script creates an empty folder structure for various temporary files and the final outputs to be saved in that is referred back to by the subsequent commands.

3)
Write all of the filepaths for all of the patient/sample BAM files into a single text file in the project directory entitled "patient_bams_list.txt", with one filepath per row, so that each row corresponds to a different BAM file.
This file is used to point the following scripts to the correct BAM files using a single number.

4)
-Run mpileup_read_count.R (must be run multiple times - once for each patient/chromosome combination)
Pass the patient/sample number (i.e. the number of the corresponding row in the patient filepath list in (3)) as the first argument and the chromosome number as the second argument.
Edit the first 11 lines of this file to match your system before running, as noted in the comments.
This script counts the numbers of reads in the BAM file for all alleles (A, C, G, T) in a specific chromosome and creates separate outputs for each, which are used in the next step.

5)
-Run get_af.R (must be run multiple times - once for each patient/chromosome combination)
Pass the patient/sample number (i.e. the number of the corresponding row in the patient filepath list in (3)) as the first argument and the chromosome number as the second argument.
Edit the first 11 lines of this file to match your system before running, as noted in the comments.
This script uses the output of mpileup_read_count.R to fill empty chromosomal matrices with allelic frequencies for all allelic reads (A, C, G, T), for each patient/chromosome combination.

6)
-Run get_mean_af.sh (must be run multiple times - once for each autosomal chromosome being examined)
Pass filepath to project directory as the first argument, and chromosome number as the second argument.
This script outputs the first half of the IncDB: mean (aggregate) allelic fractions which form the IncDB along with corresponding SD values.

7)
-Run get_sd_af.sh (must be run multiple times - once for each autosomal chromosome being examined)
Pass filepath to project directory as the first argument, and chromosome number as the second argument.
This script outputs the second half of the IncDB: standard deviations of allelic fractions which form the IncDB along with corresponding mean allelic fraction values.

OUTPUT:
The IncDB files are labelled meanAF.txt and sdAF.txt respectively and are located in the folder corresponding to the chromosome examined. E.g. chr21/meanAF.txt and chr21/sdAF.txt together form the IncDB for chromosome 21.
